# Distributed Subnetworks of Depression Defined by Direct Intracranial Neurophysiology

**DOI:** 10.1101/2020.02.14.943118

**Authors:** KW Scangos, AN Khambhati, PM Daly, LW Owen, JR Manning, JB Ambrose, E Austin, HE Dawes, AD Krystal, EF Chang

## Abstract

Quantitative biological substrates of depression remain elusive. We carried out this study to determine whether application of a novel computational approach to high spatiotemporal resolution direct neural recordings may unlock the functional organization and coordinated activity patterns of depression networks. We identified two subnetworks conserved across the majority of individuals studied. The first was characterized by left temporal lobe hypoconnectivity and pathological beta activity. The second was characterized by a hypoactive, but hyperconnected left frontal cortex. These findings identify distributed circuit activity associated with depression, link neural activity with functional connectivity profiles, and inform strategies for personalized targeted intervention.

## Introduction

Major depressive disorder (MDD) is a common, highly disabling and potentially deadly disorder that affects more than 264 million individuals worldwide (*1*). Despite significant neuroscientific advances, the biological substrate of depression remains poorly understood and new approaches that facilitate our understanding are critical. The majority of early studies seeking to characterize depression pathophysiology examined specific brain regions (ex. subgenual anterior cingulate cortex (*2-4*)), cognitive networks (ex. default mode network (*5-9*)), or univariate electrophysiological markers (ex. alpha asymmetry (*10-15*)). Yet, there is increasing evidence that depression is characterized by distributed network dysfunction beyond a single brain region or network (*16-18*).

Recent computational advancements within a network neuroscience framework have enabled researchers to model brain activity with the scope and complexity necessary to understand such distributed processes (*19*). However, detailed investigations of both the functional organization and coordinated activity patterns of depression networks have been limited by the capabilities of current imaging and electroencephalography (EEG) technologies, both indirect measures of neural activity that require a trade-off between spatial and temporal resolution. Intracranial EEG (iEEG), typically collected in patients with epilepsy for the purpose of seizure localization, has the advantage of high temporal resolution, and provides direct recordings from both cortical and subcortical brain structures. Patients with epilepsy have high rates of co-morbid depression (*20-25*) that shares origin (*26-30*) and treatment response (*31*) characteristics with primary depression. However, owing to heterogenous electrode placement across individuals, previous iEEG studies have been limited to low patient numbers and region-based approaches (*32-34*).

We hypothesized that we could apply a novel computational approach to a large unique dataset of multi-region, multi-day iEEG recordings in 54 participants to uncover distributed cortico-subcortical networks in depression. To tackle inconsistent network sampling across individuals, we utilized a method called SuperEEG (*35*) that uses the correlational structure of brain activity across the population to create a model of multiregional iEEG activity for each individual despite heterogeneous electrode placement. This model provided a highly detailed representation of brain activity across space and time and allowed us to chart out the inherent organization of the brain into functional networks. Once a generalized map of functional brain network organization was established, we quantified the multi-dimensional nature of corresponding brain dynamics to discover how rhythmic activity riding atop these functional networks differed in depressed and non-depressed individuals (*36*). Because depression has a variable presentation, we further examined how depression-associated circuitry varied across individuals in the depressed group.

We found that depression circuitry was highly distributed across cortical and subcortical structures with global dysfunction in both connectivity and spectral activity. Two unique depression subnetworks present in 89% of depressed subjects were identified. One pattern was marked by decreased connectivity across the occipitotemporal region and dominant beta band activity. The second was characterized by excessive frontal cortical connectivity with decreased activity in the alpha spectral frequency band.

## Results

### Overall Approach

Our total population consisted of 54 patients with intracranial electrodes placed for the purpose of pre-surgical mapping for treatment-resistant epilepsy. The number of electrodes per subject ranged from 33–201 (mean=120, SD=37). Our overall approach consisted of two steps – a model building step where we identified large-scale functional networks across iEEG electrodes, and a model utilization step where we related the architecture and intrinsic neural activity of functional networks to depression status (**Fig. 1**).

**Fig. 1.**
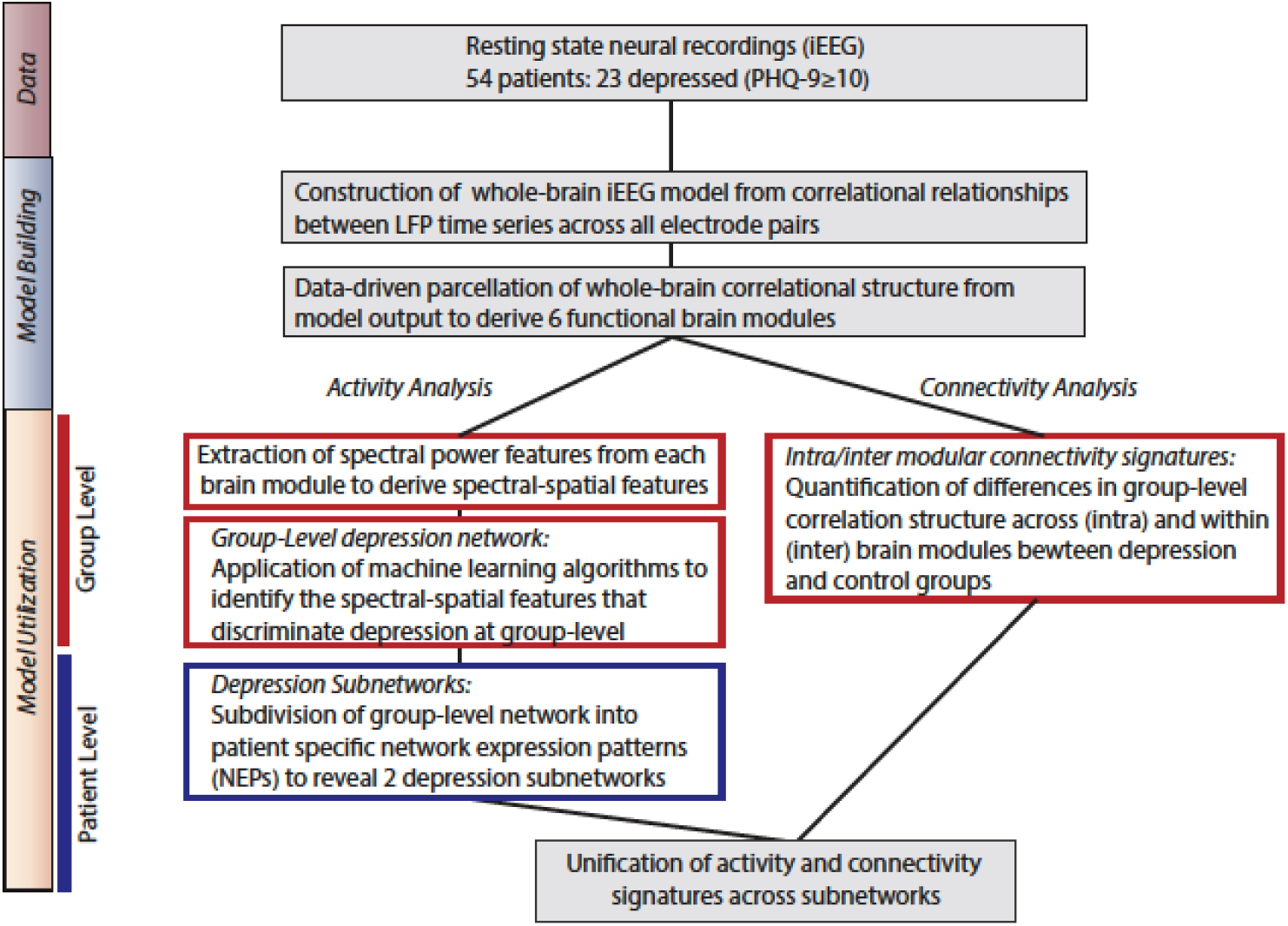
Overall approach. *Model Building:* We utilized direct neural recordings from 54 patients to construct a whole-brain model of iEEG activity based on correlational relationships of neural LFP time series signals across all electrode pairs. We then parcellated this model into functional network modules using graph theory metrics. *Model Utilization:* We used the whole-brain iEEG model to study how brain activity and connectivity measures relate to depression status. We first defined spectral power features across network modules and applied supervised machine learning to identify a group-level network features of depression (*Activity analysis*). In parallel, we identified alterations in functional network connectivity and organization between depressed and control groups (*Connectivity analysis*). Common group-level network features expressed at the individual level were clustered to identify two distinct patterns of altered activity and connectivity.

### Derivation of functional modules

We utilized a method called SuperEEG (*35*) to map continuous iEEG recordings from different patients into a common neural space and create a whole-brain, multi-subject model of iEEG recordings that served as the basis for studying distributed depression circuitry (Fig.2 A-E). This method provided an important advance over previous iEEG studies (*32-34*) that were limited to region-based analyses conducted in small samples due to heterogeneous electrode placement. In brief, to generate this model, we first constructed subject-level full-brain correlational models.

**Fig. 2.**
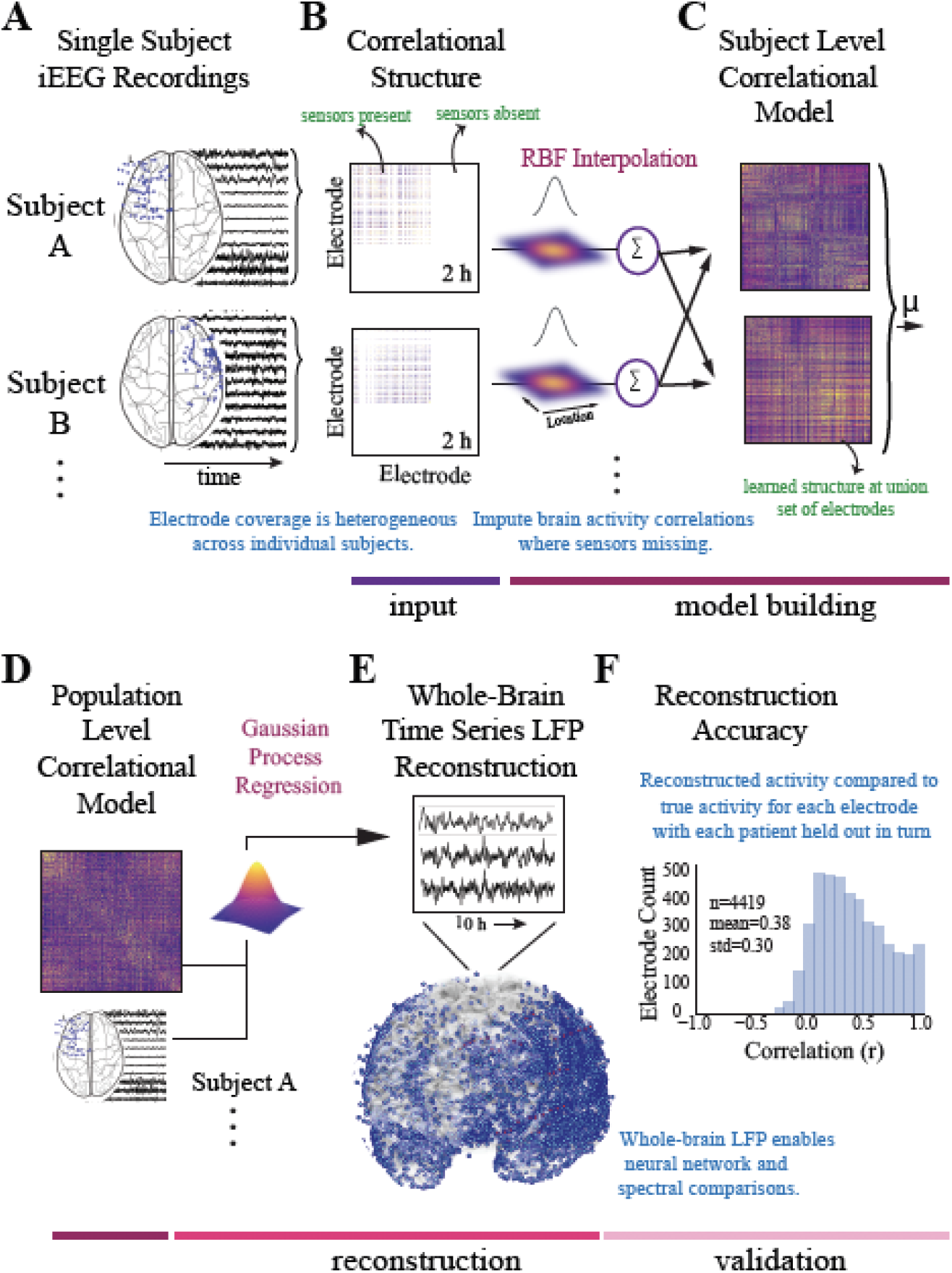
Construction of whole-brain model. **A**. To generate a multi-subject whole-brain model of iEEG activity, patient’s electrode locations across participants were first represented in a common space (Montreal Neurologic Institute (MNI) space). Electrode locations and sample recordings for a few example patients are shown. Activity was then randomly sampled in 1 min intervals across daytime hours to obtain a stable representation of brain activity across a 2-hour period. **B**. Individual inter-electrode correlation matrices were constructed for each participant at locations where electrodes were present. **C**. Subject-level full brain correlational models were then predicted using radial basis function (RBF)-weighted averages to estimate brain activity correlations at locations where sensors were not present. **D**. Subject-level correlational models were then averaged to generate a population level whole-brain correlational model. **E**. Local field potential activity for each of the 4,244 electrodes was then reconstructed using Gaussian process regression with the population-level model as a prior and activity where electrodes were present as the marginal likelihood. **F**. The distribution of the electrode signal reconstruction accuracy across our model. To obtain this distribution we built models with 55/56 patients, and then applied the model to the held-out patient, holding out each patient in turn. Correlation of the true and reconstructed signals were compared for each held-out electrode. Significance was assessed by averaging the patient level Fisher transformed correlation coefficients and comparing the distribution across patients to 0 using a t-test (t=13.94, p = 1.04e^-25^).

Interelectrode correlation matrices were constructed from activity where sensors were present and learned radial-basis function weighted averages were used to generate correlational information at locations where sensors were not present. The subject-level models were then averaged to generate a population-level model. We then used Gaussian process regression based on the population-level model and individual time series for each subject to reconstruct whole-brain local field potentials for each subject. The output of this SuperEEG (*35*) model is therefore an estimate of iEEG time series for each patient for a union set of electrodes across our total population (4,244 electrodes/per patient). Quantification of the algorithms performance has been performed previously on two large independent iEEG datasets using leave-one-out cross-validation where full-brain correlation matrices were estimated for each patient in turn using data from all the other patients (*37*). Correlation of the true and reconstructed signals were compared for each held-out electrode from the held-out patients. By using only other patients’ data to estimate activity for each held-out electrode, volume conductance or other sources of “leakage” were minimized resulting in a conservative estimate of reconstruction accuracy. Using the same approach, we found the distribution of correlations was again centered well above 0 (mean r =0.38) suggesting the algorithm estimates activity patterns substantially better than chance (**Fig.2F**). To calculate significance, we averaged the patient level fisher transformed correlation coefficients and compared the distribution across patients to 0 using a t-test (t=13.94, p = 1.04e^-25^).

After construction of the model, we next utilized principles of graph theory to identify data-driven functional networks (modules) across it. Our rationale was that the model had learned statistically correlated fluctuations between iEEG signals, akin to functional connectivity, and that a network-based approach could enhance discovery of depression circuitry over a univariate, single-region approach. We applied a modularity optimization technique known as community detection which groups electrodes into non-overlapping modules by their correlational relationships (*38, 39*) and has been used to reveal system-level disruptions in disease states (*40-46*) including MDD (*47*) (**Fig. 3**). Previous work on module detection (*48*) demonstrated that tuning a resolution parameter (*gamma)* is key to identifying modules at different topological scales of a network. In line with previous efforts that have related iEEG network structure to brain parcellations based on anatomy (*49*), we computed a similarity index between the division of electrodes into modules and the division of electrodes into anatomical structures as defined by the Lausanne atlas (*50*) for a range of resolution parameters and selected the most parsimonious match between modules and anatomical structures (*gamma*=1.19) (**Fig. S1**). We principally observed that our iEEG model was optimally parcellated into 6 stable modules (Jaccard index, p<0.05, permutation test)) and that these modules were spatially distributed and spanned multiple anatomical structures. A graph of the network and its subdivision into modules is shown in **Fig. 3B**, where module membership is indicated by the color of nodes (iEEG electrodes) and edges (inter-electrode correlation from whole-brain model). To name each module, we identified regions (hubs) with the most influential connectivity profile for each module by calculating the participation coefficient metric (*51-53*) (see *Methods*) yielding the left dorsolateral prefrontal cortical (L-DLPFC), left occipitotemporal (L-OT), left orbitofrontal cortical (L-OFC), right frontotemporal (R-FT), right medial frontal (R-MF), and mid-hemispheric modules. **Fig. 3C** shows hub locations by their mean Montreal Neurological Institute (MNI) (*50*) coordinates and associated Brodmann Areas.

**Fig. 3.**
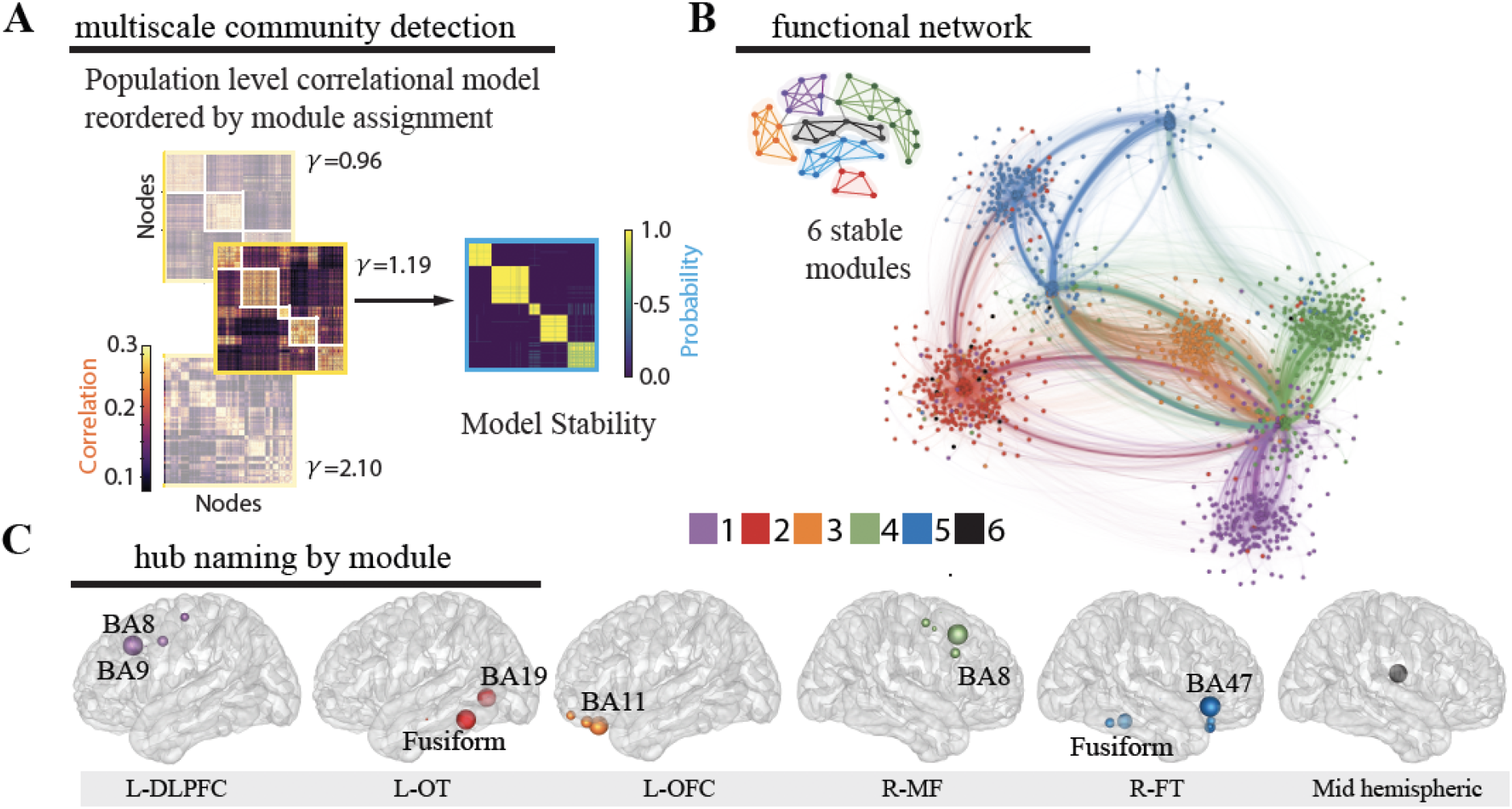
Identification of functional modules. **A**. Multiscale community detection was applied to the whole-brain model to group electrodes (nodes) into non-overlapping modules (communities) by their correlational relationships (*38, 39*). First, the population-level correlational model was reordered by the module assignment according to the modularity cost function. Network modules were identified at different levels of granularity by varying the tuning parameter, (*47, 54*). Increasing partitions the brain into increasing numbers of modules with a limit equal to the number of electrodes, as shown here for 3 values of (left). Next, the stability of this clustering at each value of was assessed by calculating module allegiance, which describes the probability that any two electrodes occupy the same module on repeated module detection (*55*) (right). A - value of 1.19 was selected by comparing the similarity of partitions generated by values of with those of a commonly used brain atlas (*50*), resulting in 6 modules. Of note, one of the modules is small and difficult to resolve in the figure. **B**. The graph of the large-scale network with module membership delineated by the color of the nodes and edges for selected γ of 1.19 is shown along with a schematic representation of the 6 modules. **C**. Hubs for each module were identified by selecting electrodes with the lowest 10% participation coefficients. Values were then averaged for each Lausanne brain region per module and weighted by the distribution of electrodes across Lausanne regions in all modules. Hub weight is indicated by the size of hub, and module assignment is indicated by hub color. Module 5 contained insufficient number of electrodes for hub identification (0.3% of total sample) and coefficients across all electrodes were utilized to name this module. L-DLPFC=left dorsolateral prefrontal cortex, L-OT=left occipitotemporal cortex, L-OFC=left orbitofrontal cortex, R-MF=right medial frontal cortex, R-FT=right frontotemporal cortex

### Relationship of functional network identification to depression status

We next investigated the whole-brain iEEG model to study brain networks underlying depression. All patients in this study were undergoing surgical mapping with multi-channel iEEG for seizure localization as part of their standard medical care. Depression was measured during the hospital stay, using the validated PHQ-9 measure (*56-58*). In accordance with literature-derived rates of depression in this population (*20-25*), 43% of our population had self-reported depression as defined by this measure (PHQ-9≥10, n=23), and 33% had mild or no symptoms of depression, which defined our control group (PHQ-9≤5, n=18). The two groups did not vary in age, sex, type of epilepsy, antidepressant usage, or anti-epileptic drug class (t-test, X^2^, p>0.4, **Table S1**).

Work over the past 50 years has attempted to utilize scalp EEG to uncover neurophysiological characteristics of depression outside of epilepsy (*59-62*). Disruption in frontal alpha power is one of the earliest and most-studied findings (*10-15*), but its value toward understanding depression is limited due to inconsistent findings across studies (*63-65*). We hypothesized that by leveraging the high temporal resolution of iEEG, as well as the direct access to subcortical structures, we could overcome limitations of scalp recordings. We calculated relative spectral power from the reconstructed time series in six frequency bands and averaged the resulting power values across time independently per module. This process yielded 36 features per participant, where each feature contained information about a spectral power band across one functional module; we therefore collectively referred to these as “spectral-spatial features.”

We then utilized a standard leave-one-out cross validated machine learning pipeline (PCA followed by logistic regression) (*66*) to identify network activity that was predictive of depression **(Fig. 4A)**. First, in line with prior iEEG analyses, we used principal component analysis (PCA) to identify a low-dimensional feature representation of spectrally band-limited neural activity across electrodes that potentially span different modules. It is important to note that while PCA and network module detection reduce the complexity and inherent collinearity in the datatset (*33, 34, 67-69*), they also reflect two non-mutually exclusive properties of brain connectivity (modules) and brain activity (principal components). Specifically, modules demarcate groups of brain regions with correlated *broadband* brain activity, irrespective of the amplitude of the activity, and principal components represent additional state-dependent neural activity that is *band-specific*, such as rhythms and oscillations (*49*), and may arise from functionally important integrative connections that span between modules (*51, 70*). This line of thinking closely resembles previously reported accounts of neural co-activation dynamics (akin to principal components) spanning multiple cognitive networks (akin to network modules) that explain inter-individual differences in task performance and cognitive traits. Second, after identifying a principal component representation of cross-module spectral-spatial network features, we utilized logistic regression (with L1 regularization) to classify subjects with depression and identify features with the greatest discriminatory power. We found that a combination of four principal components had the strongest predictive ability to detect depressed from non-depressed subjects. Their loading weights represent their contribution towards likelihood of depression. We show these weights in **Fig. 4B**. Utilizing the four most discriminative components alone, we achieved a mean classification accuracy of 80.0% on the training set and 77.4% on the test set. Significance was assessed using a permutation test (p=0.002). The same classification pipeline applied to a null model obtained from randomly permuting the target class labels 1000 times and retraining the classifier with each permutation led to an accuracy of 50.0% (70.6% training set). Together, these data suggest that a parsimonious model with four principal components, which capture major sources of variance in spectral-spatial features, can detect subjects with depression from the control group significantly better than chance.

**Fig. 4.**
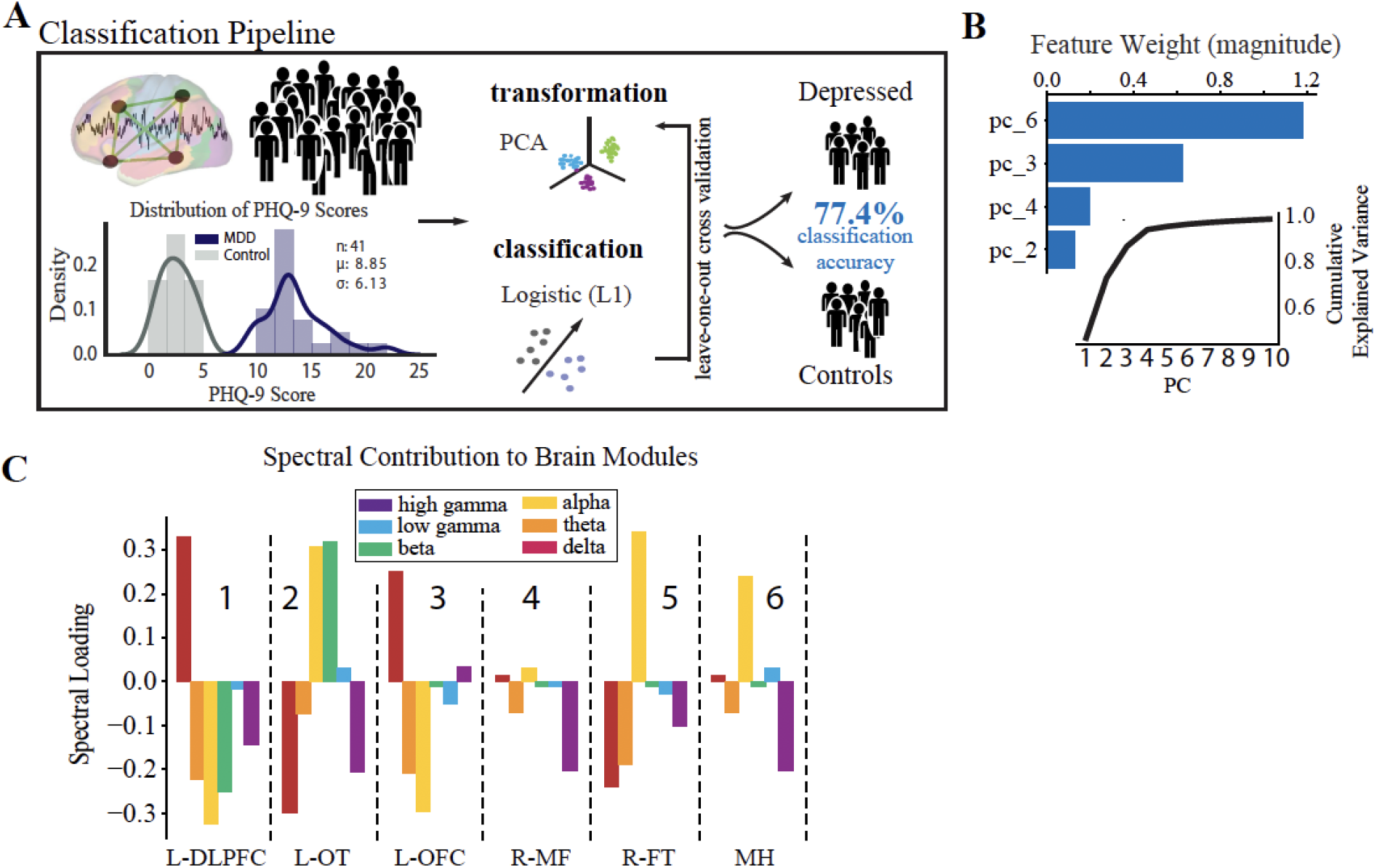
Spectral-spatial features that discriminate depression at group level. **A**. Activity analysis pipeline showing steps including power feature extraction, dimensionality reduction, transformation, and classification. The distribution of PHQ-9 scores across the depression (n=18, purple) and control groups (n=23, gray) is shown bottom left (mean PHQ-9 score 8.85, standard deviation 6.13). Power was extracted from the reconstructed time-series using the Morlet transformation in 30 s intervals across 6 frequency bands (delta = 1-4 Hz, theta = 5-8Hz, alpha = 9-12Hz, beta = 13-30Hz, low gamma (gammaL) = 31-70Hz, high gamma (gammaH) = 71-150Hz). This process yielded 25,464 spectral power features from our model (6 frequency bands x 4,244 electrodes x 2 hours). Z-scored relative power was calculated and averaged within each band across each of the 6 network modules. Power was then further averaged across time to yield 36 spectral-spatial features per participant. Principal component analysis was then used to transform the full spectral-spatial feature set, followed by logistic classification yielding 4 features that identified depression with 80.0% accuracy on the training set and 77.4% on the test set. **B**. The component weights of the four features with cumulative explained variance across the first 10 principal components shown in the inset. **C**. Spectral distribution of the 4 components was obtained by calculating the dot product between the loading weights (>0.2) for each spectral-spatial feature in the four principal components and the coefficient weighting from the classifier. Bars show the direction of change of each power band and module in relation to depression diagnosis and relate changes in spectral power associated with depression across spatially distributed brain networks. These spectral-spatial features represent the circuit activity that distinguishes depression in our population. DLPFC = dorsolateral prefrontal cortex, OT = occipitotemporal, OFC = orbitofrontal cortex, MF = medialfrontal, FT = frontotemporal, MH = mid-hemispheric

As our primary goal was to uncover the underlying biology of depression, we next turned to an examination of the individual spectral-spatial features contained within the four components. These features comprise the circuit activity that distinguishes depression in our population (for full component loadings see **Table S4**). To better interpret the biological meaning of this distributed network activity in terms of recognized brain regions (*50*) and our similarly scaled network modules, we spatially projected the four components back onto the brain (**Fig. 4C**). Specifically, we computed the dot product between the loading weights (>0.2) for each spectral-spatial feature and the coefficient weighting from the classifier. Performing this operation enabled us to show the direction of change of each power band and module in relation to depression diagnosis. It also enabled us to relate changes in spectral power associated with depression across spatially distributed brain networks. In fact, on visual inspection two gross patterns of spectral activity across the modules emerged. The first was high alpha power across the L-OT, R-FT, and mid-hemispheric modules (attention and default mode regions, modules 2,5, and 6 in **Fig. 4C**). The second was high delta and low alpha and theta power in the L-DLPFC and OFC modules (executive and limbic regions, modules 1 and 3 in **Fig. 4C**). These results suggest that low- and mid-frequency activity across broad networks characterize depression at the group level. These observations motivated the subsequent statistical analysis to define the two patterns quantitatively.

### Distinct Network Expression Patterns Define Depression

As depression is a complex disorder with heterogeneous symptom profiles across individuals, we anticipated that there would also be inter-individual heterogeneity in expression of the group-level depression network features. To assess this possibility, we first identified clusters of subjects who expressed relevant neurophysiological activity patterns by quantifying the log-odds impact on classification of each previously identified spectral-spatial feature for each subject. We next tested the distribution of feature impact on depression classification probability across participants using an agglomerative hierarchical clustering algorithm (see *Methods*) (*71-74*). We found two distinct subnetwork activity patterns (network expression patterns (NEPs)) that strongly impacted depression and subdivided our depressed population into two groups (**Fig. 5A**). The first subnetwork (NEP1) was marked by increased beta power in the L-OT module, and increased alpha and decreased delta power over the L-OT and R-FT modules. The second subnetwork (NEP2) was marked by decreased theta in the L-DLPFC, L-OFC, and R-FT modules, and decreased alpha, beta power together with increased delta power within the L-DLPFC and L-OFC modules. The presence of two subnetworks importantly demonstrated that different core features were relevant in different subjects. We next quantified the impact of each NEP on each participant’s probability of being classified as depressed using a sensitivity analysis. The difference in classification probability when one NEP was left out was attributed to the presence of that NEP, which quantified the relative importance of that NEP toward the classification of depression. **Fig. 5B** shows the probability contribution of each NEP for each subject in the depressed group (top plot) and control group (bottom plot). While we anticipated that each individual would exhibit several NEPs with differing contributions to their depression classification, an alternate pattern emerged from the data. We found that increased activity in either NEP was correlated with depression, but that each patient exhibited activity in only one of the two NEPs. Thus, depressed participants fell into two groupings based on NEP activity.

**Fig. 5.**
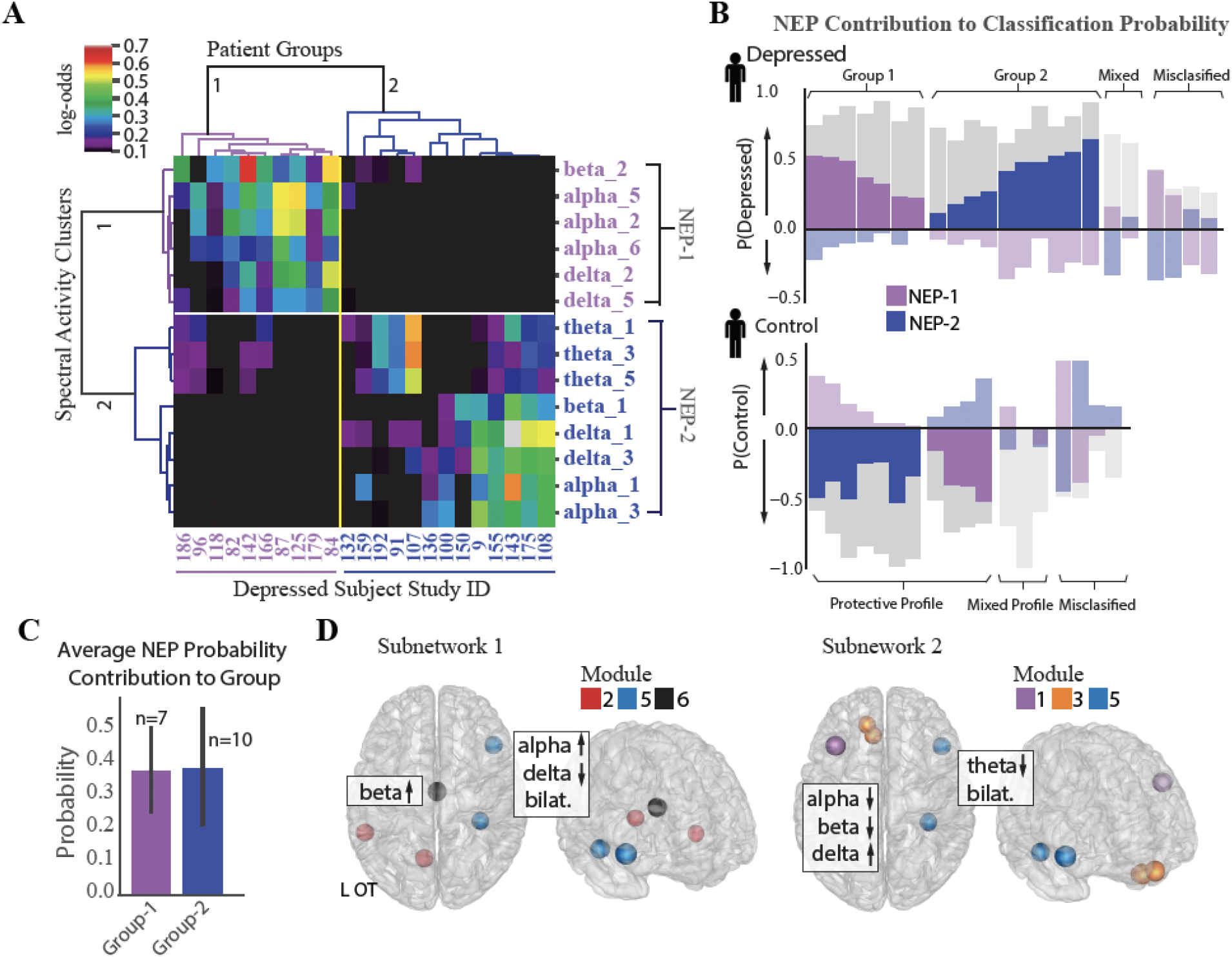
Identification of two depression subnetworks. **A**. Hierarchical clustering on log-odds of spectral-spatial features at the individual patient level showing 2 patient groups (horizontal groupings) and 2 network expression patterns (NEPs) (vertical groupings). Columns represent individual patients with patient study number shown at bottom, and rows represent spectral power across one frequency band and module (ex. alpha_1=alpha power across module 1). Magnitude of log-odds represented by color of corresponding boxes (color-bar legend top right). Spectral-spatial features associated with NEP-1 represented in purple text and those associated with NEP-2 represented in blue text. **B**. NEP probability contribution for the depressed group (top plot) and control group (bottom plot) derived from a sensitivity analysis where the probability of depression for each individual was calculated in total and with a perturbation where each NEP was held out. The probability difference was attributed to the presence of the NEP. This probability contribution is represented by the colored bars overlaid over each patient’s total probability of being depressed as derived from the machine learning classification model (gray bars, probability > 0.5 leads to classification of depression). The perturbations do not sum to produce the total classification probability; rather each quantifies the relative importance of that NEP toward depression. Bars in the positive direction indicates a positive contribution toward depression, and those in the negative direction indicate a protective contribution toward depression. Subjects where one of the two NEPs did not drive classification probability are shown in muted colors (mixed profile). Subjects classified incorrectly shown on far right of each plot (misclassified). **C**. Mean probability contribution of each NEP to two patient groups is shown. NEP-1 (purple bars) contributed most strongly to the probability of depression in the first group (mean=38% probability contribution, SD=0.13) and NEP-2 (blue bars) contributed most strongly to a second group (mean=39% probability contribution, SE=0.18). Number of participants who exhibit each NEP shown above each bar. Error bar = standard deviation. d. Direction of activity and spatial distribution of activity changes within NEP shown on glass-brain in several orientations. Hubs for each module within the NEP are designated by hub color.

Classification for the first group (37% depressed subjects) was largely driven by NEP1 (n=7, mean probability contribution =0.38, SD=0.13) alongside usually modest opposing contributions form NEP2, while classification for the second group (53% depressed subjects) was largely driven by NEP2 (n=10, mean probability contribution=0.39, SD=0.18, **Fig. 5C**), alongside more modest opposing contributions from NEP1. Classification of the remaining 11% of participants was either driven by mixed effects of both NEPs or there was little contribution from either NEP and may be evidence of additional subnetworks that were not resolved in our dataset. Two distinct groups also emerged from the control participants with NEP activity contributing here as well, but with distinct contribution profiles compared to the depressed participants. Classification for the first group (21% control patients) was driven either by mixed effects of both NEPs or little contribution of either NEP, as we anticipated. Classification for the second group, was driven by one of the two NEPs with a more modest contribution of the opposing NEP (79% of control group). We might speculate that relative NEP activity could represent either risky or conversely, protective activity profiles, and that NEP activity could be modulated in either direction to treat depression. The anatomical distribution of the two depression subnetworks is shown in **Fig. 5D**.

### Network Organization is Disrupted Across Depression Subnetworks

In addition to alterations in the spectral content of network activity in depression, previous studies have observed distinct deficiencies in connectivity across depression networks (*75-79*). A fundamental interest in neuroscience is the relationship between the brain’s neural activity and its underlying functional and structural connectivity, which remains unknown. We expected that alterations in functional network topology would be present in our depressed population and that we could delineate new relationships between activity and functional connectivity with our high-resolution dataset to more comprehensively characterize depression subnetworks. We performed a connectivity analysis using correlation of local field potential activity across modules as an estimate of functional connectivity between electrodes. The graph of our whole-brain iEEG model (**Fig. 3B**) defines correlational relationships between electrodes across our total population. We thus examined these correlational relationships across control and depressed groups independently in order to measure the relative differences of functional network organization between the two groups. Specifically, we examined the relative difference in connectivity strength across electrodes within and between functional modules. **Fig. 6A** shows the two-dimensional representation of the functional network structure for control (left) and depressed (right) groups. In comparison to the control group, we qualitatively observed an overall reduction in the segregation between modules in the depression network.

**Fig. 6.**
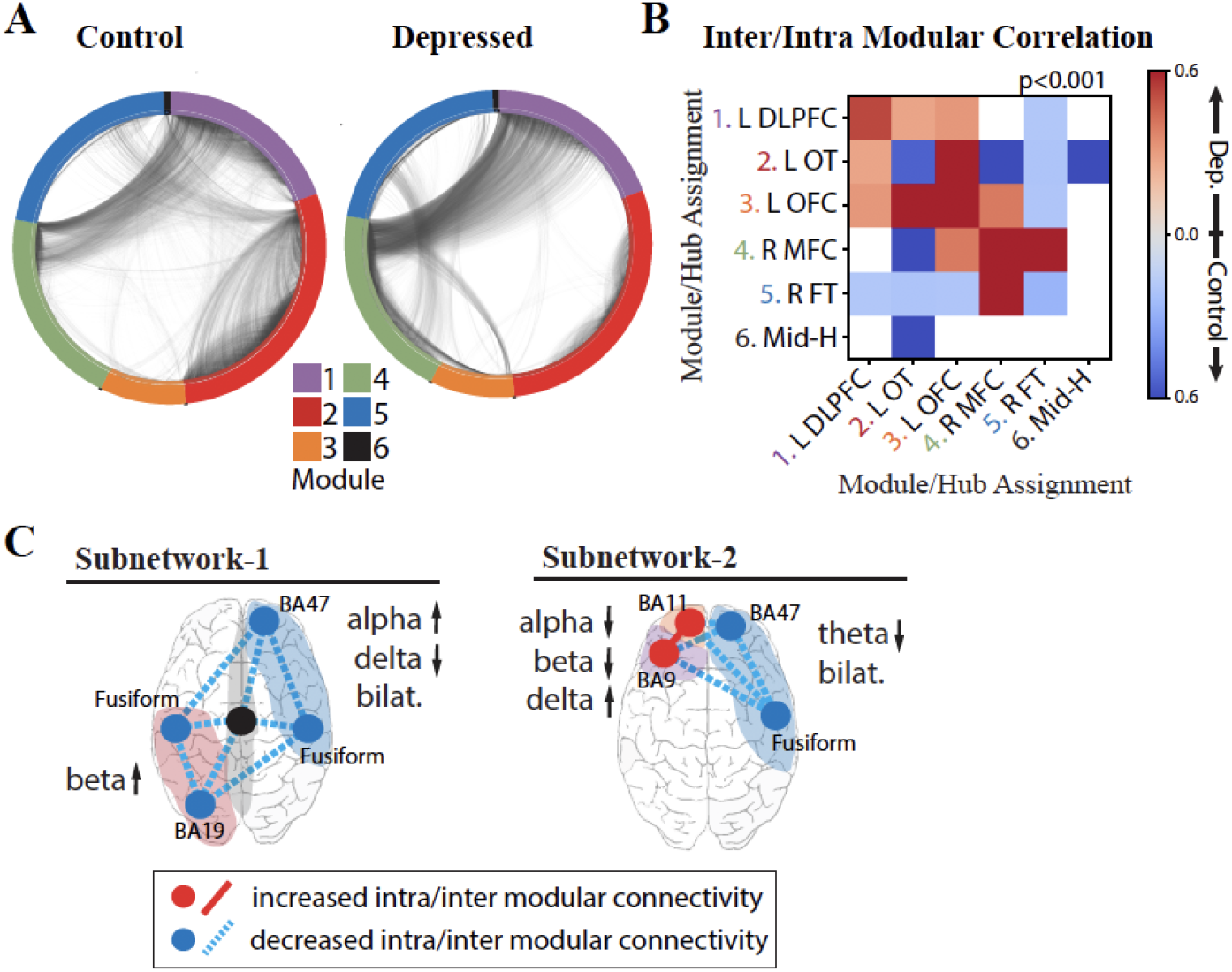
Intra- and Inter-modular connectivity signatures of depression and control groups. **A**. Connectivity structure derived using whole-brain iEEG model recalculated for the control group (left) and depressed group (right) with module membership delineated by the color of the nodes (electrodes), and edges (connections between electrodes) delineated by the black interconnecting lines. **B**. Heatmap of significant Cohen’s d values calculated from the distribution of correlation strengths between depressed and control groups for each possible module pair and compared to Cohen’s d values for a null distribution derived from permuted nodal module assignment. Those that survived multiple comparison testing (p<0.001) were retained (red: increased connectivity for depressed group; blue: increased connectivity for control group; white: not significant). **C**. Schematic of NEP-1 (left) and NEP-2 (right) showing both connectivity and spectral power underlying each pattern. Increased connectivity strength shown in red, and decreased connectivity shown in blue (hub=intramodular, line=intermodular connectivity). Color of shaded area refers to module number as shown in color legend in panel a.

To quantify these differences, we first calculated the inter- and intra-modular connectivity strength by calculating the correlation between electrodes located in the same module (intra-modular) and across different modules (inter-modular). We next compared the distribution of correlation strengths between depressed and control groups for each possible module pair using a Cohen’s *d* effect size metric. We assessed significance by retaining the Cohen’s d values that survived multiple comparisons testing with a null distribution of Cohen’s d values generated by permuting nodal module assignments (p<0.001). This analysis tests whether the effect of connectivity differences between groups is a network-wide characteristic of the depressed brain or whether the effect is localizable to specific modules. **Fig. 6B** shows the heatmap of significant Cohen’s *d* values, where a greater effect of connectivity for the depressed group is indicated in red, and lower effect of connectivity for the depressed group is indicated in blue. The results demonstrate strong evidence that, indeed, there are module-specific differences in the effect of connectivity between depressed and non-depressed individuals suggesting that modules may express hyperconnectivity or hypoconnectivity in depression depending on their anatomical localization in the brain. In the depressed group, there was overall greater frontal connectivity and weaker cross-hemispheric connectivity. Specifically, we observed greater intra-modular connectivity within L-DLPFC, L-OFC, and R-MFC modules, weaker intra-modular connectivity within L-OT and R-FT modules, and greater inter-modular connectivity between L-DLPFC, L-OFC and L-OT modules. Hubs in the insula, amygdala, temporal pole and fusiform gyrus drove the cross-module connectivity (top 10% participation coefficient, see *Methods*). We also observed a decrease in cross hemispheric connectivity in the depressed group compared to the control group (L-DLPFC/L-OFC to R-FT modules, and L-OT to R-FT/R-MFC modules), with hubs in the insula, temporal-parietal region and amygdala responsible for this decreased connectivity. The L-OFC module showed greater connectivity with the R-MFC module, and R-MFC module exhibited stronger connectivity with the R-FT module.

On the basis of the above analyses we were able to parse specific connectivity components that characterize the two depression subnetworks (**Fig. 6C**), unifying both activity and connectivity analyses across cortical and deep structures with a level of specificity that has not previously been possible. In the first subnetwork characterized by NEP1 we observed increased beta power in the L-OT module, and right-left asymmetry in the alpha and delta bands over right frontal/L-OT modules with weaker intra- and inter-modular connectivity throughout. In the second subnetwork characterized by NEP2 we observed a hyperactive left frontal cortex that was more highly connected within itself but more weakly connected to R-FT module. Lower theta bilaterally was observed in this subnetwork.

## Discussion

In this report, we present a large study of direct neural recordings aimed at identifying depression networks, made possible by multi-day iEEG recordings paired with a depression measure. The opportunity to directly record semi-chronically from cortical and subcortical structures in this manner enabled us to estimate whole-brain neural activity and incorporate both activity and connectivity analyses to resolve new subnetworks underlying depression. We found that depression is associated with a complex distributed pattern of network activity and two distinct depression subnetworks were expressed in 89% of depressed patients. These included a poorly connected occipitotemporal network characterized by heightened beta activity, and a hyperconnected frontal cortical subnetwork characterized by low alpha and theta power.

Our ability to delineate the functional organization and spectral activity patterns of depression networks with high spatiotemporal resolution relied on the application of a network neuroscience framework to the output of the SuperEEG model. Recently, Betzel and colleagues successfully applied a similar correlational network model to multi-subject iEEG recordings, followed by community detection, and found network organization to be representative of that obtained from DTI and fMRI (*49*). We further extended these findings, by applying the iEEG model to the study of disease status for the first time. The two depression subnetworks we identified are supported by previous fMRI and EEG studies of depression that have found individual components of the subnetworks in different studies including disruptions in frontal theta, temporal beta (*59*), and alpha asymmetry (*10, 11, 80-82*), decreased connectivity in the occipital, temporal, and right medial frontal regions (*16*) and higher frontal connectivity in depression (*5, 6, 83-86*). Our findings of two dichotomously expressed subnetworks may provide a partial explanation for the inconsistent findings across prior EEG studies that have predominantly focused on single frequency band or brain regions and have lacked rigorous cross-validation as noted by a recent meta-analysis (*87*).

Prior analyses of neuropsychiatric-related iEEG features have been made using components of the patient dataset used in this study (*32-34*). These efforts (*33, 34*) have focused on studying a broad emotion state rather than depression and took region-based approaches using low subject numbers due to the problem of heterogenous electrode coverage across individuals. The computational approach developed here was motivated by limitations of this prior work, enabling us to incorporate parallel information from all of our subjects despite differing electrode coverage, perform group level analysis of depression, and uncover distributed circuit activity. While our aim was to capture network dysfunction associated with depression, the two distinct ways in which activity within the NEP networks combinatorically relates to disease classification also suggest the possibility of their reflecting depression biotypes. Other studies have identified biological characteristics potentially distinguishing depression subpopulations. In one example, Drysdale and colleagues matched patient symptom profiles with fMRI connectivity features and identified several depression biotypes (*71*), one of which was dominated by globally decreased right OFC and left temporal-occipital connectivity - consistent with NEP1, and a second which exhibited increased frontal and decreased temporal connectivity - consistent with NEP2. Despite methodological differences, the similarity of these respective pairs of subnetworks – each developed in an empirical, data-driven approach using different modalities – is striking and potentially represents an emerging point of cross-scale convergence in our understanding of depression biology. Furthermore, in complement to the fMRI study, the temporal and spatial resolution of our physiology data and our network neuroscience framework for analysis has enabled a clarified picture of the patterns of modular and distributed neural activity that may underlie the expression of depression subtypes in different patients. Deeper exploration of these putative biotypes await further study.

Functional connectivity informs longer time-scale organization of neural populations whereas functional activity informs moment-to-moment behavior of neural populations. Our finding that some brain regions show distinct changes in both activity and connectivity, while other regions, such as the right medial frontal region (module 4), demonstrate connectivity differences alone suggest that depression is both a state-invariant connectivity disorder and a state-dependent activity disorder. This relationship might explain why traditional antidepressant medications can take 6-8 wks to start working, yet ketamine can improve symptoms on the same day of administration (*88*). It is possible that the presence of aberrant activity over long periods of time could shape network connectivity via plasticity or that changed connectivity patterns can impact the timing and flow of normal neural activity. Future work using high temporal resolution iEEG could inform how symptom-states and depression traits are integrated at the level of distributed neural circuits.

We acknowledge some weaknesses in the results presented. Depression in epilepsy is thought to arise from similar origins to primary depression (ex. stress (*26*), inflammation (*27*), circuit dysfunction (*30*)), and is responsive to antidepressants (*31*) suggesting it can provide valuable insight into depression more broadly. It remains unknown whether the depression networks we identified are related to the presence of epilepsy. Our categorical approach using the PHQ-9 to identify depressed patients was straightforward to apply in the context of complex data and has direct clinical relevance. However, it also selects inherently imperfect diagnostic boundaries and limited our capacity to examine variation in depression among subjects. Furthermore, as this was a cross-sectional investigation, some patients in the control group had a history of depression treated with ongoing antidepressant use but were not depressed per the PHQ-9 at the time of the study. Future analyses could explore how neural signatures vary with symptom severity in addition to alternative dimensional approaches which have the potential benefit of mapping neural features onto symptom profiles (*71, 72*).

While our whole-brain iEEG model was extensive in coverage, we did not have electrodes placed in all brain regions, including some regions implicated in depression (*89-93*) and the density of electrode sampling varied across brain regions leading to uncertainty in the accuracy of estimation in sparsely sampled areas (*94*). We dealt with this constraint by discounting the effect of each individual node degree before running community detection and comparing network measures to a null model that accounted for overall node density. Furthermore, our prior work has shown no reliable correlation between reconstruction accuracy and density (*35*).

SuperEEG relies on accurate reconstruction of held-out activity patterns. While accuracy of this algorithm is significantly above chance and similar to the test-retest reliability of fMRI in redetecting estimated activity (*95*), improved reconstruction is an important area for future work. With advancements in data processing capabilities and accessibility we may be able to reduce assumptions and the estimation burden, extend coverage to more brain regions, and utilize larger samples. Indeed, work to integrate our findings with network features from high spatial resolution MRI is already underway by our group. Finally, while ideally we would have independent test and training datasets for the machine learning used for classification, we utilized leave-one-out cross validation due to our sample size.

In light of methodological limitations of the whole-brain iEEG modelling approach it is important that we highlight a critical, yet subtle, conclusion regarding the saliency of biological signatures in background iEEG recordings that are associated with depression. Indeed, the SuperEEG approach reconstructs just a portion of the verum iEEG signal – the remaining unexplained portion may stem from subject-specific variation in connectivity (*96, 97*), state-dependent variability in connectivity (*98, 99*) within subjects, or statistical noise. It follows that of this faithfully reconstructed portion of the iEEG signal, we found that higher-order principal components of spectral-spatial iEEG activity were most important for identifying patients with depression. Taken together, we speculate that depression may in fact have a low-dimensional network representation that is widely pervasive in the iEEG signal but represents just a small portion of iEEG signal dynamics. Importantly, we found that alternate ML pipelines converged on these same low-dimensional features and the composite feature set resembles neural features of depression that have been cited in previous neuroimaging studies. Thus, there is high likelihood that the neural features we have found reflect circuit physiology that is stereotyped to depression.

Through the current study, we identified two novel subnetworks of depression. The results have important implications for disease subtyping, diagnosis, treatment planning, and monitoring of depression status. These subnetworks could form the basis for interventions at many different potential control points along each subnetwork and suggest that interventions that change both connectivity and spectral power could be promising. For example, they provide a mechanistic rationale for practitioner’s choice between right and left DLPFC vs. OFC targets for repetitive transcranial magnetic stimulation (*71, 100*). Evidence of high activity in one network pattern, countered by an anti-weighting of the other pattern further suggests the existence of protective or high-risk profiles and the possibility of preventative treatments. A library of new treatment targets and frequency-specific treatment parameters (*101, 102*) could enable a new wave of closed-loop interventional therapies that personalize treatment based on neurophysiological signals.

## Materials and Methods

### Patient Characterization

Participants included 54 adults (49% female) aged 20-67 who had medication-refractory epilepsy and were undergoing intracranial monitoring as part of their standard medical care. Neural data from these participants comprised our full dataset and was utilized to build the whole-brain iEEG model of LFP time-series. Participants were screened for depression following electrode implantation and concurrent with neural recordings using the Patient Health Questionnaire-9 (PHQ-9), a 9-item self-report instrument validated for depression screening (*56-58*). A score ≥ 10 defined the MDD group (moderate depression) and a score ≤ 5 defined the non-depressed control group generating a sample of 23 depressed subjects (56%) and 18 controls (44%). The remaining 13 patients were used in the first step of the study (model *building*) but not the second (model *utilization*). Data comprised a consecutive series of patients recruited from University of California, San Francisco and Kaiser Permanente, Redwood City, California over a 5-year period. This study was approved by the University of California, San Francisco Institutional Review Board with written informed consent provided by all subjects.

Patients’ antiepileptic medications (AEDs) were withdrawn as part of standard clinical care. However, to control for possible effects of medication on neural activity in the depressed and control groups we examined the number of patients in each group that were on AEDs associated with depression(*103*) using a chi squared test. Depressive disorders have been shown to be significantly increased with AEDs that have strong GABA-ergic properties such as barbituates, tiagabine, vigabatrin, and topiramate (*29*). Zonisamide (*104*) and perampanel (*105*) also have been shown to be associated with depression. There was no significant difference in the number of patients on such medications in the control and depressed group (**Table S1**) providing evidence that medication usage is not driving the neurophysiological group differences.

### Electrode Implantation and Localization

Subdural grid, strip, and depth electrodes (AdTech, Racine, WI; or Integra, Plainsboro, NJ) were implanted using standard neurosurgical techniques. Subjects underwent pre-operative 3 Tesla brain magnetic resonance imaging (MRI) and post-operative computed tomography (CT) scan to localize electrodes in patient-centered coordinates using an open source python package for preprocessing imaging data for use in iEEG recordings (*106*). The steps included warping brain reconstructions to a common Montreal Neurologic Institute (MNI) template and merging electrode locations across subjects. Surface warpings were then generated by projecting pial surfaces of the subject and template brains into a spherical coordinate space and aligning the surfaces in that space. Depth warping was then performed using a combination of volumetric and surface warping (*107*). For visualization, pre-operative T1-weighted MRI scans were pre-registered with the post-operative CT using Statistical Parametric Mapping software SPM12 and pial surface 3D reconstructions were generated using FreeSurfer (*108*).

### Data Acquisition and Pre-Processing

Data acquisition of iEEG recordings were acquired using the Natus EEG clinical recording system at a sampling rate of 1-2 kHz. Standard iEEG/ECoG pre-processing techniques were conducted in python including application of a 2-250Hz bandpass filter, notch filters at line noise frequency and harmonics (60Hz, 120Hz, 180Hz, 240Hz), down sampling to 512Hz, and common average referencing to the mean of all channels. The data were acquired across a range of behaviors while the patient was in the epilepsy monitoring unit.

### Construction of Whole-Brain iEEG Model

A functional connectivity imputation technique was utilized to estimate whole-brain iEEG activity for each subject (SuperEEG). This method involved four main steps outlined in **Fig. 2**. First, pre-processed iEEG signals were chunked into 60s non-overlapping blocks and filtered for putative epileptiform activity or artifacts. We then randomly sampled the 60s intervals across daytime hours (8am-10pm) and concatenated them into 2 hours blocks, each representative of naturalistic activity. To filter the activity of putative epileptiform activity or artifacts we identified signals that deviated from the norm using kurtosis, a measure of infrequent extreme peaked deviations (*109*). Individual 60s blocks were retained if the number of artifactual signals (kurtosis>10) was less than 5% of the total sample. The same kurtosis threshold was then applied to each 2h block to further eliminate noisy electrodes. The kurtosis threshold of 10 was tested in Owen et al. 2020 on a similar intracranial dataset and verified by eye on a subset of samples by a provider trained in Clinical Neurophysiology. Second, the iEEG functional connectivity model was built by averaging patient level connectivity models. The patient level model leveraged known inter-electrode correlations to estimate temporal correlations at locations that were not present. This was accomplished by using the fact that the strength of relationships between sensors are proportional to neural proximity. As the distance between two sensors increases the correlation weakens. The rate of falloff was modeled using a gaussian radial basis function (*rbf*) at locations *η* of each known patient electrode to estimate the unknown correlations between *x* and *η*.

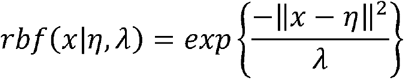

where *λ* was set to 20 based on prior work that sought to maximize both spatial resolution and generalizability to any brain location (*51*). This relationship was utilized as the Weight matrix (*w*) and applied to every electrode set (*i,j*) for each patient as

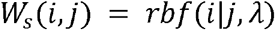

where the rbf function expands the single patient to include estimated temporal recording relationships from sensors in the full population. Third, the Weight matrix was then combined with the single patient correlation matrix (*C*_*s*_(*i,j*) to estimate each patient’s full population anatomical connection structure using,

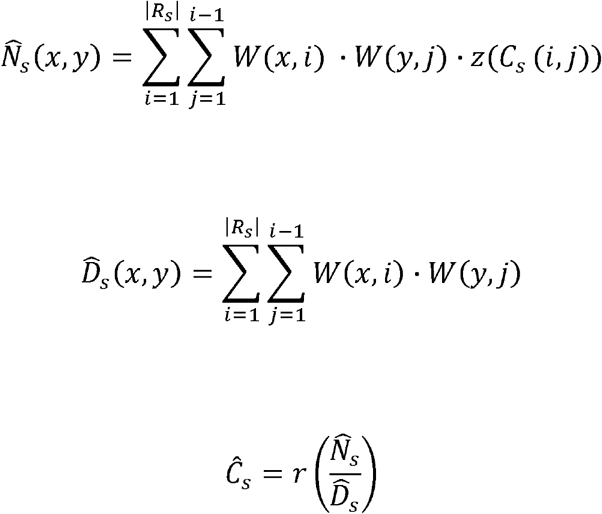

where r(·) represents the Fisher-z transformation. The patient specific estimates of full brain correlation matrix 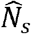 and the spatial proximity weighting matrix 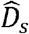 were combined in a weighted average to compute a single whole-brain expected correlation matrix 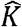 as

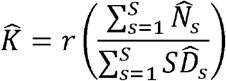

Finally, estimated LFP activity for each electrode was reconstructed using Gaussian process regression according to

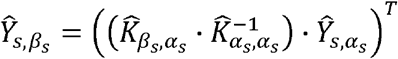

where *s* represents the patient, *α*_*s*_ is the set of indices where we had observed recordings for patient *s* and *β*_*s*_ is the set of indices that were to be estimated. The submatrix 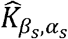 represents the estimated correlations between locations of known activity and locations of reconstructed activity. The submatrix 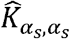 represents correlations between known neural recordings.

Construction of the above model required extensive computational resources. Therefore, we sought to utilize the minimum required information required to obtain the majority of information and enable computational feasibility. Using the 10h benchmark as the largest feasible model we could build, we compared 2, 4, 6 and 8 hour models to the 10 hour model and found that the difference in adding additional time beyond 2h was marginal and could be computed at a fraction of the computational cost. We therefore utilized the 2h model for further analysis (**Fig. S1A**).

Validation of this method on two independent datasets has been performed previously, demonstrating reconstruction accuracies for held out electrodes are significantly better than random guessing (*35*). We performed this same validation on electrodes from our dataset. To do so, a model with all but one patient was constructed and then applied to the held-out patient. The correlations of the true and reconstructed signals were compared for each electrode. The distribution of reconstruction accuracies is shown in **Fig. 2F**. Significance was assessed by averaging the patient level fisher transformed correlation coefficients and comparing the distribution across patients to 0 using a t-test.

### Signal Processing

Standard signal processing techniques were applied to the time-series activity across all reconstructed electrodes. This included continuous wavelet transformation using the Morlet transform wavelet method (6-cycles) (*110*) performed in 30s intervals to obtain power spectra in 6 frequency bands (delta = 1-4Hz, theta = 5-8Hz, alpha = 9-12Hz, beta = 13-30Hz, low gamma (gammaL) = 31-70Hz, high gamma (gammaH) = 71-150Hz). Relative power was calculated by dividing the power of each frequency band by the total power across the 6 frequency bands for each electrode. Signals were summarized by taking the mean power across time for each spectral band and were z-scored across patients.

### Electrode Clustering into Functional Modules

We used a well-validated technique, multiscale community detection, to identify distributed functional sub-networks (modules) across the model (*38, 39, 48, 111-113*). The purpose of module detection was to uncover the inherent organization of the brain’s correlational structure from the output of the SuperEEG model. We conceptualize a network module as a distinct property of connectivity organization, akin to validated atlas parcellations (*50*) but specifically designed for functional rather than structural data. Atlases apply boundaries to brain regions based on structural or functional organization derived from coarse-scale neuroimaging and thus, while they provide a useful validation for our data-driven parcellation scheme, there is no reason to assume their boundaries will perfectly align with neural signals at the millimeter scale of iEEG.

Individual functional connectivity models generated in the whole-brain iEEG reconstruction were used as a starting point in this analysis. Using the Louvain algorithm (*114*), we identified an optimal parcellation of electrodes into discrete functional modules by maximizing a modularity cost function defined by the following relationships,

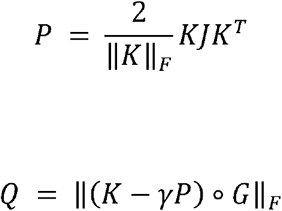

where *J* is a ones matrix, ° is the Hadamard product and Gi,j is 0 if node pair (*i,j*) are assigned to different modules and 1 if the pair is assigned to the same module, *Q* is modularity, *K* is the connection weights (correlation) between node *i* and *j, P* is the Newman-Girvan null model (*39*) and *γ* is the weighting of that null model which is tuned to obtain network modules of different sizes. As the network fractures into many small modules for higher *γ*, the overall modularity *Q* decreases. We examined network modularity at values of *γ*, between 0.5 and 2.1. We first assessed the stability of clustering at each value of *γ*, by examining module allegiance (*55*), calculated by repeating module detection 100 times and evaluating the probability that two electrodes occupied the same module. We then selected an optimal value of *γ*, by assessing how the iEEG modules co-localized with brain structures derived from the 234 anatomically distinct brain areas defined by Cammoun et al. (2012) (*50*) referred to as the Lausanne atlas. In accordance with prior work (*49*), we computed a similarity index (Rand score index (*115*)) between the division of electrodes into modules and the division of electrodes into anatomical structures for the range of resolution parameters (**Fig. S1B**). Significance was assessed by a permutation test where the null model was generated by randomly assigning electrodes to each module and calculating the confidence interval of the similarity index generated from 1,000 random permutations and tested at significance level 0.05 for a 2-tailed test. Two similarity peaks were identified, with values of γ that generated 6 and 1,855 modules respectively. The peak with the highest modularity (lowest number of clusters) was selected for further analysis due to our goal of examining the brain at a low level of granularity. This selection enabled subsequent classification of activity across these clusters without overfitting our model. While we report our results based on this most parsimonious match between modules and anatomical structures (*γ* =1.19), we verified that the assignment of electrodes into slightly coarser and slightly finer modules (1 < *γ*, < 2.1) did not substantially alter our ability to predict subjects with depression (**Fig. S1B, *red***). Finally, we assessed the distribution of electrodes that were assigned to each module across the main anatomical regions defined by *Cammoun et al. (2012) (50)* (**Fig. S1**).

### Assigning Names to Modules

We assigned a name to each module by examining the location of each module’s most influential electrodes. We utilized the participation coefficient (PaC), which is a degree-based measure of network connectivity that describes a node’s functional interaction within and across network modules (*51-53*). This metric is typically utilized to identify influential hubs across a large-scale network. We utilized it in our study to identify the location of hubs that were most important for driving connectivity in each module identified through community detection. Groups of electrodes with low PaC values indicate hubs with high intramodular connectivity, also known as provincial hubs (*116*). Similarly, connector hubs are those with high PaCs and drive intermodular connectivity. The PaC describes the weight of edges from node i to all other nodes in the same module relative to the weight of edges from that node to all nodes in the network according to

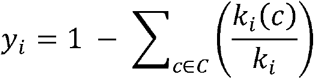

where y_i_ is node *i*’s participation coefficient, C is the set of all modules, *k*_*i*_(*c*)is the sum of all correlations between node *i* and other members of module C and *k*_*i*_ is the sum of all correlations between node *i* and members of all modules. We calculated the PaC for each electrode across our model, and then selected those with high and low participation values (top/bottom 10%). We then grouped these selected nodes by Lausanne atlas region, eliminating or combining a minority of regions due to having too few electrodes for analysis. We addressed the non-uniform distribution of electrodes across the model by then assigning each Lausanne region a score according to the following hub weight:

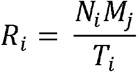

where *N*_*i*_ is the number of selected electrodes (top/bottom 10%) in Lausanne region *i*, *M*_*j*_ is the number of selected electrodes in Lausanne region *i* of module *j*, and *T*_*j*_ is the number of total electrodes across modules in Lausanne region *i*. Hubs were those Lausanne regions with the highest hub weight. Hub location was identified by averaging the MNI coordinates of electrodes within each hub. The full list of Lausanne regions and hub weights is shown in **Fig. S2 and Tables S2-3**. The purpose of the identified hubs in the present report was primarily descriptive and helped relate the computational model to known brain regions and structure; all subsequent analyses utilized the population set of electrodes across the full model.

### Classification

We utilized a machine learning algorithm validated with leave-one-out cross validation to identify distributed neural circuit features that discriminated depression. We first averaged local field potentials across the electrodes within each module and then decomposed the signals into common spectral band to identify 36 features (6 frequency bands x 6 modules) that contained information both about the spectral power and the location of the activity. These features, referred to as spectral-spatial features, served as our starting feature space for entry into our classification pipeline. Transformation with principal component analysis (PCA) (*117*) followed by methods for feature selection and subsequent discrimination have been used on previous iEEG classification problems (*33, 34*). We followed a similar pipeline. PCA enabled us to reduce collinearity and identify latent variables that described maximally distinct network-based components inclusive of multiple frequency bands and spatial modules. A logistic regression classifier was then utilized to classify depression based on accuracy using a sparsity promoting regulizer (L1) which selects fewer principal components while controlling for overfitting. PCA and logistic classification were performed within the cross-validation loop where a model is trained on all subjects but one, and then tested on the held-out subject with each subject held-out in turn. We report mean accuracy (balanced to group-size) across the cross-validation iterations. To further asses our model validity, we repeated our classification pipeline on a null model obtained from randomly permuting the target class labels 1000 times and used a permutation test to assess significance between the true and null model accuracy distributions.

In order to control for possible differences in epileptiform activity residual to data-cleaning across the modules we calculated line-length, a commonly utilized measure for the detection of epileptiform activity (*118*) for each electrode and averaged across the modules. We used a logistic regression model to show that line-length across the six modules was not a significant predictor of depression status (R^2^=0.15, p=0.13, logistic regression).

### Hierarchical clustering to identify depression networks

We reasoned that we could utilize the group-level network to identify common features that defined depression at the individual level. To do so, we mapped the principal component values (feature loadings ≥0.2) back to the original feature space weighted by the logistic regression coefficients. This provided the log-odds impact of each original feature. We then tested the distribution of feature impact on depression classification probability across depressed participants by grouping similar log-odds impact covariates (thresholded at 0.15) utilizing an agglomerative hierarchical clustering algorithm (*71-74*). A log-odds threshold of 0.15 was selected because it retained classification results for 98% of subjects while isolating the most contributory spectral-spatial features (see **Fig. S3A** for non-thresholded model for comparison). The clustering yielded both patient and feature groupings that defined neurophysiological network expression patterns (NEPs) of depression. We quantified the impact of these NEPs on each participant’s probability of being classified as depressed by performing a sensitivity analysis where we withheld each NEP and then attributed the probability decrement from the total classification probability to the withheld activity pattern. Two patient groupings were defined based on the relative contribution of each NEP to classification probability. Subjects in whom the contribution was very modest or mixed across the NEPs were placed into a third mixed group. We also ran this analysis on the boundary patients who had mild symptoms of depression but did not reach threshold (PHQ-9<10) for depression (**Figure S3B**).

### Connectivity Analysis

In order to understand the effect of depression on network topology we examined changes in connectivity across the control and depressed subjects. First, inter- and intramodular connectivity strengths were assessed by looking at the correlations between all electrodes within the same module (intramodular) and the correlations between electrodes across all pairs of modules (intermodular). Next, to assess whether the effect of connectivity differences between groups is a network-wide characteristic of the depressed brain or whether the effect is localizable to specific modules, we used a Cohen’s d effect size metric and compared the distribution of correlation strengths across depressed and control groups for each possible module pair. To assess significance across these connections we generated a null distribution of Cohen’s d values for each module pair and retained the true Cohen’s d values that survived multiple comparisons testing (*p*<0.001).

## Acknowledgements

We thank the members of the Chang Lab for assistance.

## Funding

This work was supported by National Institutes of Health award K23NS110962. (K.W.S), NARSAD Young Investigator grant from the Brain and Behavioral Research Foundation (K.W.S), and a Ray and Dagmar Dolby Family Fund through the Department of Psychiatry at the University of California, San Francisco (K.W.S, A.D.K.) and a Brain Initiative grant (SUBNETS, E.F.C.). Dr Chang receives research support from National Institutes of Health, New York Stem Cell Foundation, the Howard Hughes Medical Institute, the McKnight Foundation, the Shurl and Kay Curci Foundation, and the William K. Bowes Foundation. Dr. Krystal receives support from National Institutes of Health, PCORI, Janssen, Jazz, Axsome, Reveal Biosensors.

## Author Contributions

K.W.S., A.N.K., E.F.C., A.D.K. conceived the study. K.W.S., A.N.K., P.M.D. analyzed and interpreted the data. A.J.B and A.E. contributed to data collection. J.R.M and L.W.O contributed to data analysis methods. K.W.S wrote the manuscript with significant input from all authors. All authors reviewed and approved the manuscript.

## Competing Interests

K.W.S., A.N.K., P.M.D., J.R.M and L.W.O report no competing interests. A.K. consults for Eisai, Evecxia, Ferring, Galderma, Harmony Biosciences, Idorsia, Jazz, Janssen, Merck, Neurocrine, Pernix, Sage,and Takeda. E.F.C. has patents related to brain stimulation for neuropsychiatric conditions, brain mapping, and speech neuroprosthesis. Dr. Chang has given talks related to epilepsy treatment for Neuropace and Cyberonics/Livanova.

## Data and Materials Availability

Raw data is provided in Fig. S3 for the hierarchical clustering and in Tables S2 (participation coefficient) and 3 (principal component analysis). All datasets generated and analyzed in the current study are available from the corresponding authors upon request. All custom code, statistical analysis and visualizations were performed in Python and are demonstrated in a set of Jupyter notebooks available for download. The following open-source code was used as described in the methods: SuperEEG (https://github.com/ContextLab/supereeg).

## References

1. G. B. D. Disease, I. Injury, C. Prevalence, Global, regional, and national incidence, prevalence, and years lived with disability for 354 diseases and injuries for 195 countries and territories, 1990-2017: a systematic analysis for the Global Burden of Disease Study 2017. Lancet 392, 1789–1858 (2018).

2. K. N. Botteron, M. E. Raichle, W. C. Drevets, A. C. Heath, R. D. Todd, Volumetric reduction in left subgenual prefrontal cortex in early onset depression. Biol Psychiatry 51, 342–344 (2002).

3. S. H. Kennedy et al., Changes in regional brain glucose metabolism measured with positron emission tomography after paroxetine treatment of major depression. Am J Psychiatry 158, 899–905 (2001).

4. S. Yoshimura et al., Rostral anterior cingulate cortex activity mediates the relationship between the depressive symptoms and the medial prefrontal cortex activity. J Affect Disord 122, 76–85 (2010).

5. M. D. Greicius et al., Resting-state functional connectivity in major depression: abnormally increased contributions from subgenual cingulate cortex and thalamus. Biol Psychiatry 62, 429–437 (2007).

6. Y. I. Sheline, J. L. Price, Z. Yan, M. A. Mintun, Resting-state functional MRI in depression unmasks increased connectivity between networks via the dorsal nexus. Proc Natl Acad Sci U S A 107, 11020–11025 (2010).

7. R. Bluhm et al., Resting state default-mode network connectivity in early depression using a seed region-of-interest analysis: decreased connectivity with caudate nucleus. Psychiatry Clin Neurosci 63, 754–761 (2009).

8. S. Grimm et al., Altered negative BOLD responses in the default-mode network during emotion processing in depressed subjects. Neuropsychopharmacology 34, 932–943 (2009).

9. X. Zhu et al., Evidence of a dissociation pattern in resting-state default mode network connectivity in first-episode, treatment-naive major depression patients. Biol Psychiatry 71, 611–617 (2012).

10. I. H. Gotlib, C. Ranganath, J. P. Rosenfeld, Frontal EEG alpha asymmetry, depression, and cognitive functioning. Cognition Emotion 12, 449–478 (1998).

11. J. B. Henriques, R. J. Davidson, Left frontal hypoactivation in depression. J Abnorm Psychol 100, 535–545 (1991).

12. A. H. Kemp et al., Disorder specificity despite comorbidity: resting EEG alpha asymmetry in major depressive disorder and post-traumatic stress disorder. Biol Psychol 85, 350–354 (2010).

13. M. A. Diego, T. Field, M. Hernandez-Reif, CES-D depression scores are correlated with frontal EEG alpha asymmetry. Depress Anxiety 13, 32–37 (2001).

14. L. M. Kentgen et al., Electroencephalographic asymmetries in adolescents with major depression: influence of comorbidity with anxiety disorders. J Abnorm Psychol 109, 797–802 (2000).

15. N. Jaworska, P. Blier, W. Fusee, V. Knott, alpha Power, alpha asymmetry and anterior cingulate cortex activity in depressed males and females. J Psychiatr Res 46, 1483–1491 (2012).

16. I. M. Veer et al., Whole brain resting-state analysis reveals decreased functional connectivity in major depression. Front Syst Neurosci 4, (2010).

17. L. L. Zeng et al., Identifying major depression using whole-brain functional connectivity: a multivariate pattern analysis. Brain 135, 1498–1507 (2012).

18. F. Liu et al., Abnormal amplitude low-frequency oscillations in medication-naive, first-episode patients with major depressive disorder: a resting-state fMRI study. J Affect Disord 146, 401–406 (2013).

19. D. S. Bassett, O. Sporns, Network neuroscience. Nat Neurosci 20, 353–364 (2017).

20. E. Fuller-Thomson, S. Brennenstuhl, The association between depression and epilepsy in a nationally representative sample. Epilepsia 50, 1051–1058 (2009).

21. F. Gilliam, H. Hecimovic, Y. Sheline, Psychiatric comorbidity, health, and function in epilepsy. Epilepsy Behav 4 Suppl 4, S26–30 (2003).

22. B. P. Hermann, J. E. Jones, Intractable epilepsy and patterns of psychiatric comorbidity. Adv Neurol 97, 367–374 (2006).

23. B. P. Hermann, M. Seidenberg, B. Bell, Psychiatric comorbidity in chronic epilepsy: identification, consequences, and treatment of major depression. Epilepsia 41 Suppl 2, S31–41 (2000).

24. D. Rai et al., Epilepsy and psychiatric comorbidity: a nationally representative population-based study. Epilepsia 53, 1095–1103 (2012).

25. W. A. Swinkels, J. Kuyk, R. van Dyck, P. Spinhoven, Psychiatric comorbidity in epilepsy. Epilepsy Behav 7, 37–50 (2005).

26. A. C. Wulsin, M. B. Solomon, M. D. Privitera, S. C. Danzer, J. P. Herman, Hypothalamic-pituitary-adrenocortical axis dysfunction in epilepsy. Physiol Behav 166, 22–31 (2016).

27. A. Vezzani, J. French, T. Bartfai, T. Z. Baram, The role of inflammation in epilepsy. Nat Rev Neurol 7, 31–40 (2011).

28. M. Mula, B. Schmitz, Depression in epilepsy: mechanisms and therapeutic approach. Ther Adv Neurol Disord 2, 337–344 (2009).

29. B. Schmitz, Effects of antiepileptic drugs on mood and behavior. Epilepsia 47 Suppl 2, 28–33 (2006).

30. E. Gleichgerrcht, M. Kocher, L. Bonilha, Connectomics and graph theory analyses: Novel insights into network abnormalities in epilepsy. Epilepsia 56, 1660–1668 (2015).

31. A. M. Kanner, Depression in epilepsy: prevalence, clinical semiology, pathogenic mechanisms, and treatment. Biol Psychiatry 54, 388–398 (2003).

32. K. W. Scangos et al., Pilot Study of An Intracranial Electroencephalography Biomarker of Depressive Symptoms in Epilepsy. J Neuropsychiatry Clin Neurosci, appineuropsych19030081 (2019).

33. L. A. Kirkby et al., An Amygdala-Hippocampus Subnetwork that Encodes Variation in Human Mood. Cell 175, 1688–1700 e1614 (2018).

34. O. G. Sani et al., Mood variations decoded from multi-site intracranial human brain activity. Nat Biotechnol 36, 954–961 (2018).

35. L. L. W. Owen, Muntianu, T.A., Heusser, A.C., Daly, P., Scangos, K., Manning, J.R., A Gaussian process model of human electrocorticographic data. Cereb Cortex Forthcoming, (2020).

36. S. Gu et al., The Energy Landscape of Neurophysiological Activity Implicit in Brain Network Structure. Sci Rep 8, 2507 (2018).

37. L. L. W. Owen et al., A Gaussian Process Model of Human Electrocorticographic Data. Cereb Cortex, (2020).

38. V. D. Blondel, J. L. Guillaume, R. Lambiotte, E. Lefebvre, Fast unfolding of communities in large networks. J Stat Mech-Theory E, (2008).

39. M. E. Newman, Finding community structure in networks using the eigenvectors of matrices. Phys Rev E Stat Nonlin Soft Matter Phys 74, 036104 (2006).

40. Z. J. Chen, Y. He, P. Rosa-Neto, G. Gong, A. C. Evans, Age-related alterations in the modular organization of structural cortical network by using cortical thickness from MRI. Neuroimage 56, 235–245 (2011).

41. J. Bruno, S. M. Hosseini, S. Kesler, Altered resting state functional brain network topology in chemotherapy-treated breast cancer survivors. Neurobiol Dis 48, 329–338 (2012).

42. M. Cao, N. Shu, Q. Cao, Y. Wang, Y. He, Imaging functional and structural brain connectomics in attention-deficit/hyperactivity disorder. Mol Neurobiol 50, 1111–1123 (2014).

43. Y. Sun et al., Disrupted functional brain connectivity and its association to structural connectivity in amnestic mild cognitive impairment and Alzheimer’s disease. PLoS One 9, e96505 (2014).

44. A. F. Alexander-Bloch et al., Disrupted modularity and local connectivity of brain functional networks in childhood-onset schizophrenia. Front Syst Neurosci 4, 147 (2010).

45. Q. Yu et al., Altered topological properties of functional network connectivity in schizophrenia during resting state: a small-world brain network study. PLoS One 6, e25423 (2011).

46. C. L. Keown et al., Network organization is globally atypical in autism: A graph theory study of intrinsic functional connectivity. Biol Psychiatry Cogn Neurosci Neuroimaging 2, 66–75 (2017).

47. Y. He et al., Reconfiguration of Cortical Networks in MDD Uncovered by Multiscale Community Detection with fMRI. Cereb Cortex 28, 1383–1395 (2018).

48. D. S. Bassett et al., Robust detection of dynamic community structure in networks. Chaos 23, 013142 (2013).

49. R. F. Betzel et al., Structural, geometric and genetic factors predict interregional brain connectivity patterns probed by electrocorticography. Nat Biomed Eng, (2019).

50. L. Cammoun et al., Mapping the human connectome at multiple scales with diffusion spectrum MRI. J Neurosci Meth 203, 386–397 (2012).

51. M. A. Bertolero, B. T. Yeo,M. D’Esposito, The modular and integrative functional architecture of the human brain. Proc Natl Acad Sci U S A 112, E6798–6807 (2015).

52. R. Guimera, L. A. Amaral, Cartography of complex networks: modules and universal roles. J Stat Mech 2005, nihpa35573 (2005).

53. M. Rubinov, O. Sporns, Complex network measures of brain connectivity: uses and interpretations. Neuroimage 52, 1059–1069 (2010).

54. J. O. Garcia, A. Ashourvan, S. F. Muldoon, J. M. Vettel, D. S. Bassett, Applications of community detection techniques to brain graphs: Algorithmic considerations and implications for neural function. Proc IEEE Inst Electr Electron Eng 106, 846–867 (2018).

55. D. S. Bassett, M. Yang, N. F. Wymbs, S. T. Grafton, Learning-induced autonomy of sensorimotor systems. Nat Neurosci 18, 744–751 (2015).

56. K. Kroenke, R. L. Spitzer, J. B. Williams, The PHQ-9: validity of a brief depression severity measure. J Gen Intern Med 16, 606–613 (2001).

57. R. L. Spitzer, K. Kroenke, J. B. Williams, Validation and utility of a self-report version of PRIME-MD: the PHQ primary care study. Primary Care Evaluation of Mental Disorders. Patient Health Questionnaire. JAMA 282, 1737–1744 (1999).

58. R. L. Spitzer, J. B. Williams, K. Kroenke, R. Hornyak, J. McMurray, Validity and utility of the PRIME-MD patient health questionnaire in assessment of 3000 obstetric-gynecologic patients: the PRIME-MD Patient Health Questionnaire Obstetrics-Gynecology Study. Am J Obstet Gynecol 183, 759–769 (2000).

59. J. J. Newson, T. C. Thiagarajan, EEG Frequency Bands in Psychiatric Disorders: A Review of Resting State Studies. Front Hum Neurosci 12, 521 (2018).

60. V. E. Pollock, L. S. Schneider, Quantitative, waking EEG research on depression. Biol Psychiatry 27, 757–780 (1990).

61. R. Thibodeau, R. S. Jorgensen, S. Kim, Depression, anxiety, and resting frontal EEG asymmetry: a meta-analytic review. J Abnorm Psychol 115, 715–729 (2006).

62. S. Olbrich, R. van Dinteren, M. Arns, Personalized Medicine: Review and Perspectives of Promising Baseline EEG Biomarkers in Major Depressive Disorder and Attention Deficit Hyperactivity Disorder. Neuropsychobiology 72, 229–240 (2015).

63. C. Gold, J. Fachner, J. Erkkila, Validity and reliability of electroencephalographic frontal alpha asymmetry and frontal midline theta as biomarkers for depression. Scand J Psychol 54, 118–126 (2013).

64. S. A. Reid, L. M. Duke, J. J. Allen, Resting frontal electroencephalographic asymmetry in depression: inconsistencies suggest the need to identify mediating factors. Psychophysiology 35, 389–404 (1998).

65. N. van der Vinne, M. A. Vollebregt, M. van Putten, M. Arns, Frontal alpha asymmetry as a diagnostic marker in depression: Fact or fiction? A meta-analysis. Neuroimage Clin 16, 79–87 (2017).

66. M. R. Arbabshirani, S. Plis, J. Sui, V. D. Calhoun, Single subject prediction of brain disorders in neuroimaging: Promises and pitfalls. Neuroimage 145, 137–165 (2017).

67. J. R. Manning, S. M. Polyn, G. H. Baltuch, B. Litt, M. J. Kahana, Oscillatory patterns in temporal lobe reveal context reinstatement during memory search. Proc Natl Acad Sci U S A 108, 12893–12897 (2011).

68. K. W. Scangos, R. D. Weiner, E. C. Coffey, A. D. Krystal, An Electrophysiological Biomarker That May Predict Treatment Response to ECT. J ECT 35, 95–102 (2019).

69. J. R. Manning, M. R. Sperling, A. Sharan, E. A. Rosenberg, M. J. Kahana, Spontaneously reactivated patterns in frontal and temporal lobe predict semantic clustering during memory search. J Neurosci 32, 8871–8878 (2012).

70. R. F. Betzel, J. D. Medaglia, D. S. Bassett, Diversity of meso-scale architecture in human and non-human connectomes. Nat Commun 9, 346 (2018).

71. A. T. Drysdale et al., Resting-state connectivity biomarkers define neurophysiological subtypes of depression. Nat Med 23, 28–38 (2017).

72. K. A. Grisanzio et al., Transdiagnostic Symptom Clusters and Associations With Brain, Behavior, and Daily Function in Mood, Anxiety, and Trauma Disorders. JAMA Psychiatry 75, 201–209 (2018).

73. E. Ravasz, A. L. Somera, D. A. Mongru, Z. N. Oltvai, A. L. Barabasi, Hierarchical organization of modularity in metabolic networks. Science 297, 1551–1555 (2002).

74. J. Rihel et al., Zebrafish behavioral profiling links drugs to biological targets and rest/wake regulation. Science 327, 348–351 (2010).

75. T. Chen et al., Anomalous single-subject based morphological cortical networks in drug-naive, first-episode major depressive disorder. Hum Brain Mapp 38, 2482–2494 (2017).

76. M. S. Korgaonkar, A. Fornito, L. M. Williams, S. M. Grieve, Abnormal structural networks characterize major depressive disorder: a connectome analysis. Biol Psychiatry 76, 567–574 (2014).

77. A. Lord, D. Horn, M. Breakspear, M. Walter, Changes in community structure of resting state functional connectivity in unipolar depression. PLoS One 7, e41282 (2012).

78. Y. Sun, S. Hu, J. Chambers, Y. Zhu, S. Tong, Graphic patterns of cortical functional connectivity of depressed patients on the basis of EEG measurements. Conf Proc IEEE Eng Med Biol Soc 2011, 1419–1422 (2011).

79. J. Zhang et al., Disrupted brain connectivity networks in drug-naive, first-episode major depressive disorder. Biol Psychiatry 70, 334–342 (2011).

80. A. J. Tomarken, R. J. Davidson, R. E. Wheeler, L. Kinney, Psychometric properties of resting anterior EEG asymmetry: temporal stability and internal consistency. Psychophysiology 29, 576–592 (1992).

81. R. E. Wheeler, R. J. Davidson, A. J. Tomarken, Frontal brain asymmetry and emotional reactivity: a biological substrate of affective style. Psychophysiology 30, 82–89 (1993).

82. J. B. Henriques, R. J. Davidson, Regional brain electrical asymmetries discriminate between previously depressed and healthy control subjects. J Abnorm Psychol 99, 22–31 (1990).

83. G. S. Alexopoulos et al., Functional connectivity in the cognitive control network and the default mode network in late-life depression. J Affect Disord 139, 56–65 (2012).

84. T. Frodl et al., Functional connectivity bias of the orbitofrontal cortex in drug-free patients with major depression. Biol Psychiatry 67, 161–167 (2010).

85. E. A. Nofzinger et al., Alterations in regional cerebral glucose metabolism across waking and non-rapid eye movement sleep in depression. Arch Gen Psychiatry 62, 387–396 (2005).

86. W. Cheng et al., Medial reward and lateral non-reward orbitofrontal cortex circuits change in opposite directions in depression. Brain 139, 3296–3309 (2016).

87. A. S. Widge et al., EEG Biomarkers for Treatment Response Prediction in Major Depressive Illness. Am J Psychiatry 176, 82 (2019).

88. A. McGirr et al., A systematic review and meta-analysis of randomized, double-blind, placebo-controlled trials of ketamine in the rapid treatment of major depressive episodes. Psychol Med 45, 693–704 (2015).

89. H. S. Mayberg et al., Cingulate function in depression: a potential predictor of treatment response. Neuroreport 8, 1057–1061 (1997).

90. P. Riva-Posse et al., A connectomic approach for subcallosal cingulate deep brain stimulation surgery: prospective targeting in treatment-resistant depression. Mol Psychiatry 23, 843–849 (2018).

91. D. A. Malone, Jr. et al., Deep brain stimulation of the ventral capsule/ventral striatum for treatment-resistant depression. Biol Psychiatry 65, 267–275 (2009).

92. W. R. Marchand et al., Aberrant functional connectivity of cortico-basal ganglia circuits in major depression. Neurosci Lett 514, 86–90 (2012).

93. C. Hamani et al., The subcallosal cingulate gyrus in the context of major depression. Biol Psychiatry 69, 301–308 (2011).

94. L. W. Owen, Heusser A.C., Manning, J.R., A Gaussian process model of human electrocorticographic data. Preprint at https://www.biorxiv.org/content/10.1101/121020v2 (2018).

95. C. M. Bennett, M. B. Miller, How reliable are the results from functional magnetic resonance imaging? Ann N Y Acad Sci 1191, 133–155 (2010).

96. S. Mueller et al., Individual variability in functional connectivity architecture of the human brain. Neuron 77, 586–595 (2013).

97. E. S. Finn et al., Functional connectome fingerprinting: identifying individuals using patterns of brain connectivity. Nat Neurosci 18, 1664–1671 (2015).

98. R. M. Hutchison, T. Womelsdorf, J. S. Gati, S. Everling, R. S. Menon, Resting-state networks show dynamic functional connectivity in awake humans and anesthetized macaques. Hum Brain Mapp 34, 2154–2177 (2013).

99. R. M. Hutchison et al., Dynamic functional connectivity: promise, issues, and interpretations. Neuroimage 80, 360–378 (2013).

100. K. Feffer et al., 1Hz rTMS of the right orbitofrontal cortex for major depression: Safety, tolerability and clinical outcomes. Eur Neuropsychopharmacol 28, 109–117 (2018).

101. L. Chanes, R. Quentin, C. Tallon-Baudry, A. Valero-Cabre, Causal frequency-specific contributions of frontal spatiotemporal patterns induced by non-invasive neurostimulation to human visual performance. J Neurosci 33, 5000–5005 (2013).

102. L. Cocchi, A. Zalesky, Personalized Transcranial Magnetic Stimulation in Psychiatry. Biol Psychiatry Cogn Neurosci Neuroimaging 3, 731–741 (2018).

103. S. Nadkarni, O. Devinsky, Psychotropic effects of antiepileptic drugs. Epilepsy Curr 5, 176–181 (2005).

104. J. R. White et al., Zonisamide discontinuation due to psychiatric and cognitive adverse events: a case-control study. Neurology 75, 513–518 (2010).

105. F. Rugg-Gunn, Adverse effects and safety profile of perampanel: a review of pooled data. Epilepsia 55 Suppl 1, 13–15 (2014).

106. L. S. Hamilton, D. L. Chang, M. B. Lee, E. F. Chang, Semi-automated Anatomical Labeling and Inter-subject Warping of High-Density Intracranial Recording Electrodes in Electrocorticography. Front Neuroinform 11, 62 (2017).

107. G. Postelnicu, L. Zollei, B. Fischl, Combined volumetric and surface registration. IEEE Trans Med Imaging 28, 508–522 (2009).

108. B. Fischl, FreeSurfer. Neuroimage 62, 774–781 (2012).

109. B. Akbarian, A. Erfanian, Automatic Seizure Detection Based on Nonlinear Dynamical Analysis of EEG Signals and Mutual Information. Basic Clin Neurosci 9, 227–240 (2018).

110. S. J. Schiff, A. Aldroubi, M. Unser, S. Sato, Fast wavelet transformation of EEG. Electroencephalogr Clin Neurophysiol 91, 442–455 (1994).

111. A. R. Antony et al., Simultaneous scalp EEG improves seizure lateralization during unilateral intracranial EEG evaluation in temporal lobe epilepsy. Seizure 64, 8–15 (2019).

112. P. J. Mucha, T. Richardson, K. Macon, M. A. Porter, J. P. Onnela, Community structure in time-dependent, multiscale, and multiplex networks. Science 328, 876–878 (2010).

113. O. Sporns, R. F. Betzel, Modular Brain Networks. Annu Rev Psychol 67, 613–640 (2016).

114. V. D. Blondel, J.-L. Guillaume, R. Lambiotte, E. Lefebvre, Fast unfolding of communities in large networks. Journal of Statistical Mechanics: Theory and Experiment 2008, P10008 (2008).

115. B. Misic et al., Network-Level Structure-Function Relationships in Human Neocortex. Cereb Cortex 26, 3285–3296 (2016).

116. M. P. van den Heuvel, O. Sporns, Network hubs in the human brain. Trends Cogn Sci 17, 683–696 (2013).

117. H. Hotelling, Analysis of a complex of statistical variables into principal components. Journal of Educational Psychology 24, 417-441, 498-520 (1933).

118. L. Guo, D. Rivero, J. Dorado, J. R. Rabunal, A. Pazos, Automatic epileptic seizure detection in EEGs based on line length feature and artificial neural networks. J Neurosci Methods 191, 101–109 (2010).

