## Supplementary Materials for "Distributed Subnetworks of Depression Defined by Direct Intracranial Neurophysiology"

**Supplementary Figures**

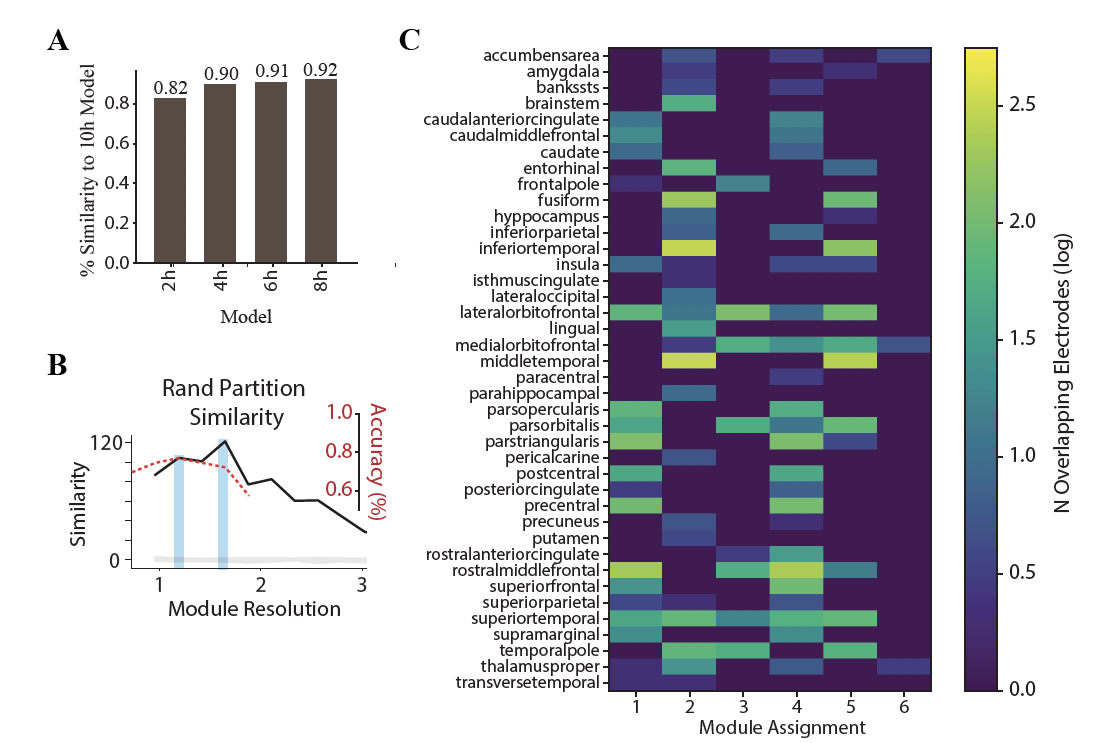

**Fig. S1. Model Assessment. A.** Comparison of 10h model to models derived from 2, 4, 6, and 8h models showing the computationally feasible 2h model contained the majority of information in a 10h model. Percent similarity to 10h model defined by the frobenius norm of the distance between this model and the 2, 4, 6, 8h models normalized by the frobenius norm of the 10h model. **B.** Multiscale community detection was utilized to identify network organization (*48*). To select a level of granularity, the module resolution parameter was varied, and the resulting network organization was compared to the 234 anatomically distinct brain areas defined by Cammoun et al. (2012) (*50*) referred to as the Lausanne atlas. Rand partition similarity scores are shown on left sided y-axis, revealing two peaks (highlighted in blue). Grey line shows results of permutation test randomly assigning electrodes to each module and calculating the confidence interval of the similarity index generated from 1000 random permutations and tested at significance level 0.05 for a 2-tailed test. Classification accuracy is shown on right sided y-axis across a range of module resolution parameter values showing that selection a different granularity does not substantially alter our ability to predict subjects with depression. **C.** Heat-map of overlap between module resolution parameter of 1.19 and the Lausanne brain atlas reflecting the joint spatial distribution of electrode assignments to modules detected through unsupervised community detection (x-axis) and to anatomical brain regions parcellated based on the Lausanne brain atlas (Cammoun et al. 2012) (*50*). Each electrode is localized to a single Lausanne brain region and is assigned a single module. Darker colors imply that fewer electrodes within a brain region belong to a particular module and brighter colors imply that more electrodes within a brain region belong to a particular module. The data demonstrate that modules are comprised of electrodes that span multiple, spatially-distributed brain regions and that the density of anatomical electrode localization is largely complimentary between modules. However, we also observed that certain brain structures appear in multiple modules due to more heterogeneous functional connectivity profiles, suggesting they express a more integrative topological function that spans multiple brain networks.

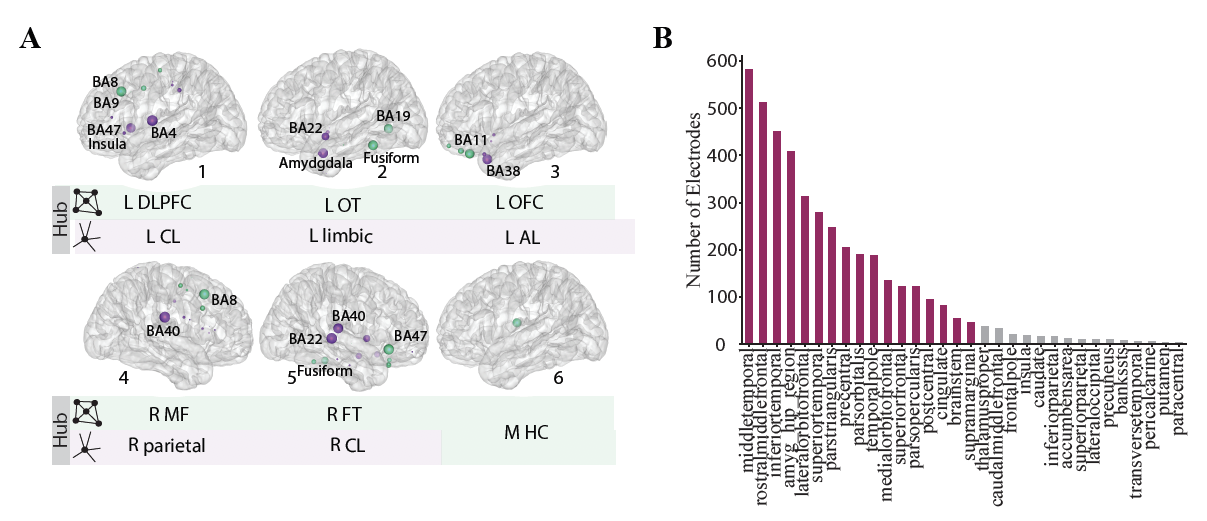

**Fig. S2. Assigning Hub Names to Modules.** **A.** Provincial (green) and connector (purple) hubs are shown on a glass brain for each of the 6 modules along with their associated Brodmann Areas. Hubs were derived from the calculation of the participation coefficient (PaC) across each module. Provincial hubs were groups of electrodes with low PaC and connector hubs were identified as those electrodes with high PaC values. The size of the hub reflects hub importance to the module, determined by the PaC value weighted by electrode density per Lausanne brain region. **B.** The Lausanne brain regions selected for the calculations in a. are shown in purple and determined by the number of electrodes per region. L DLPFC = left dorsolateral prefrontal, L OT = left occipitotemporal, L OFC = left orbitofrontal, R MF = right medial frontal, R FT = right frontotemporal, L CL = left corticolimbic, L Limbic = left limbic, L AL= left association limbic, R parietal = right parietal, R CL = right corticolimbic, M HC = mid-hemispheric cluster

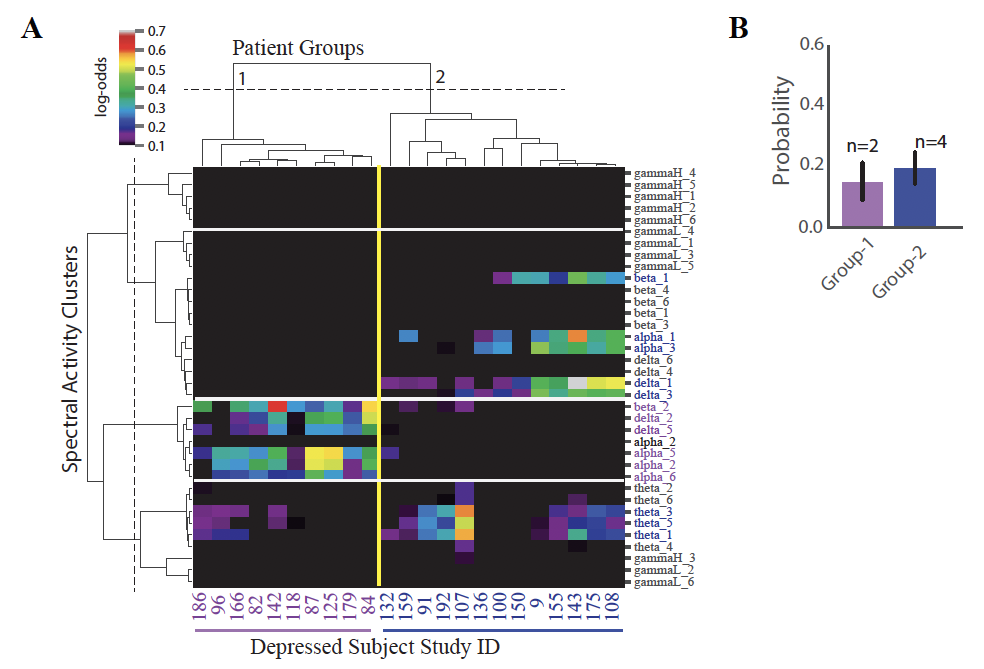

**Fig. S3. Depression Subnetwork Identification. A.** Hierarchical clustering on the individual representation of the group-level biomarker was performed with a threshold log-odds of 0.15 in order to select features that were most contributary to the classification of depression. The results of this analysis without this thresholding is shown here for comparison. As shown, the patient grouping is identical without thresholding. NEP-1 and NEP-2 again characterize Group 1 and Group 2 with NEP-2 split into two clusters. **B.** Mean probability contribution of each NEP to Group 1 and 2 for the boundary patients 5<PHQ9<10. As for the depressed population, NEP-1 (purple bars) contributed most strongly to the probability of depression in the first group (mean=15% probability contribution, SD=0.09) and NEP-2 (blue bars) contributed most strongly to a second group (mean=20% probability contribution, SE=0.06). However, as expected, the probability contribution was lower (see Fig. 5C for comparison). Number of participants who exhibit each NEP shown above each bar. Error bar = standard deviation.

**Supplementary Tables**

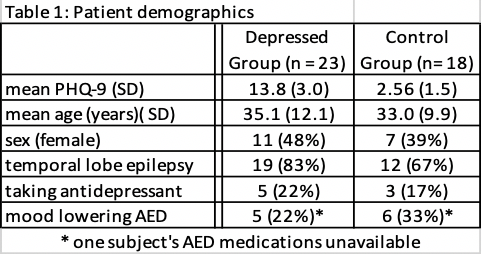

**Table S1. Patient demographics.** Mood lowering medications included barbituates, tiagabine, vigabatrin, topiramate, zonisamide, and perampanel.

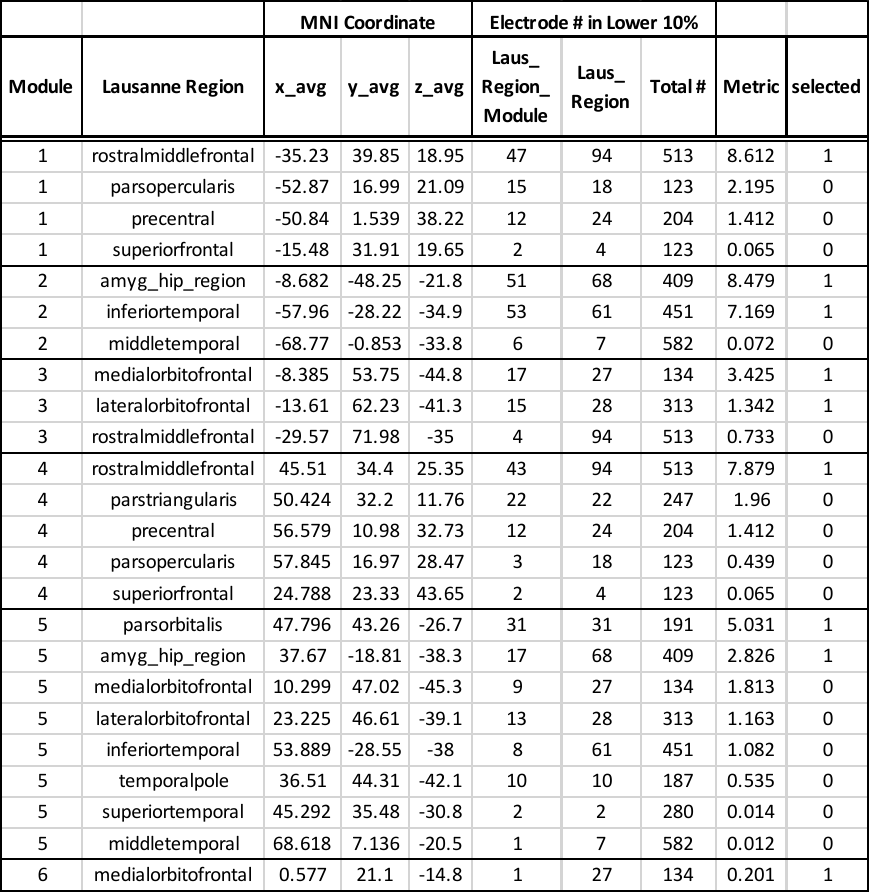

**Table S2: Hubs selected for module naming (Provincial Hubs).** First column indicates module assignment, second column indicates Lausanne region labels per module with electrodes within the bottom 10% PaC. The number of electrodes belonging to that Lausanne region within the top 10% participation coefficient indicated in column entitled (Laus_Rregion_Module), the number of electrodes belonging to that region in total is indicated by column entitled (Laus_Region), and the total number of electrodes within that Lausanne region across modules is indicated in column entitled (Total #). The metric describes the average PaC weighted by density for electrodes within each Lausanne region. The average MNI coordinate of electrodes in those regions is shown. The last column indicates whether or not that hub was selected for module naming.

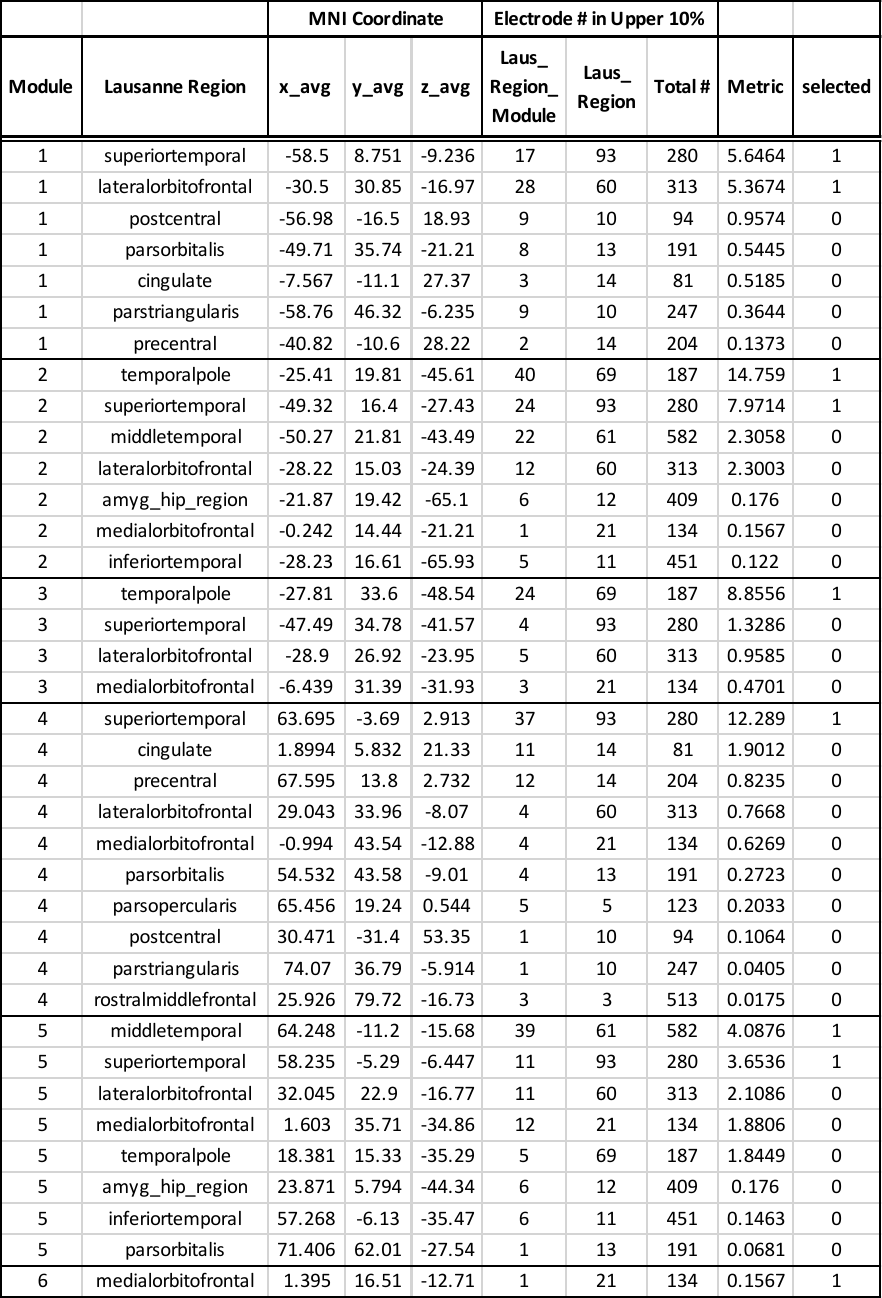
 **Table S3.** **Hubs selected for module naming (Connector Hubs).** Columns as in Table 1, for the PaC in the top 10 percentile.

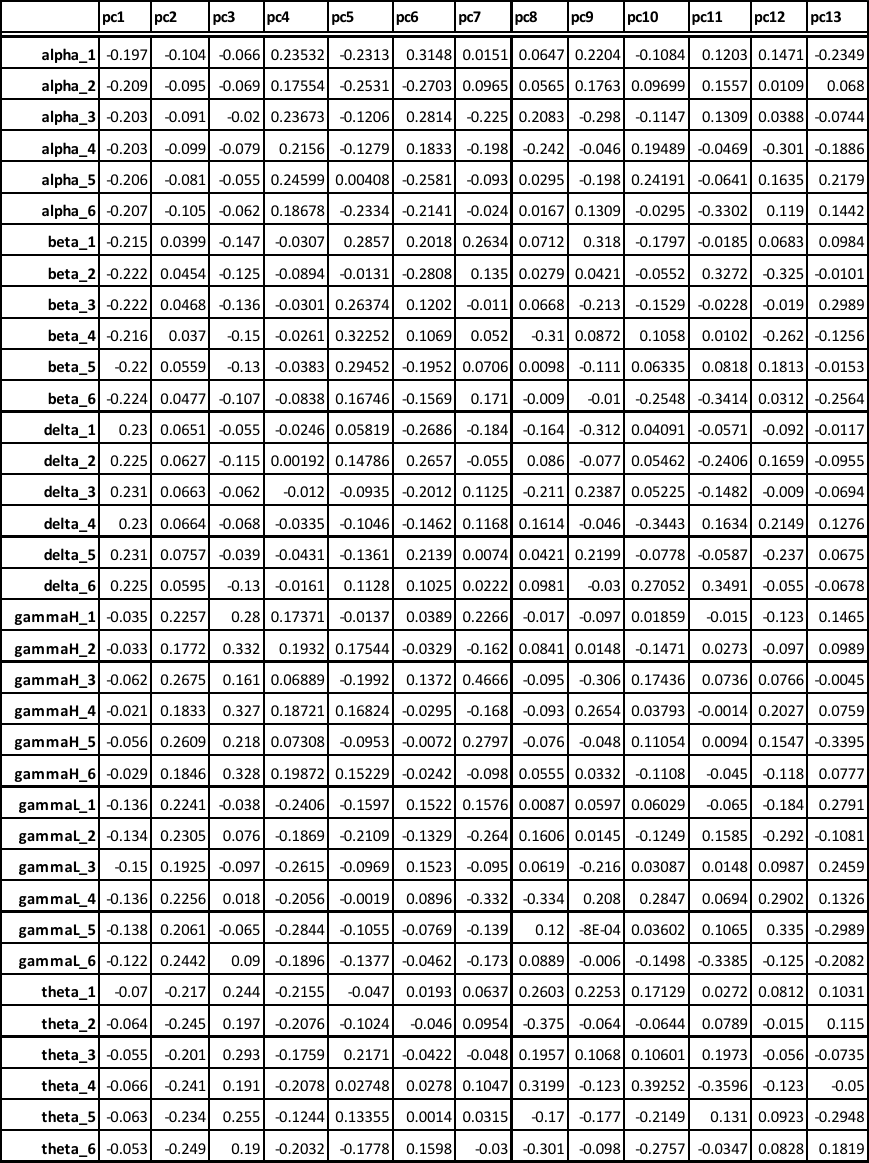

**Table S4.** Component Loadings on the individual spectral-spatial features across 13 components derived from the principal component analysis.
